# Adversity severity shapes caregiving predictors and neural correlates of infant social behavior

**DOI:** 10.64898/2026.09.30.755360

**Authors:** Allison Su, Elaina Regier, Chloe Lim, Isaac McKie, Ben Perez-Flesler, Vishal Win, Isabellah Nikitah, Brianna C. Bullock, Damaris Chacon, Lucy Sparks Mendez, Keita Fogus, Kellie L. Tamashiro, Patrese A. Robinson, Maya Opendak

**Affiliations:** Kennedy Krieger Institute, Baltimore MD 21205 USA; Solomon H. Snyder Department of Neuroscience, Johns Hopkins University School of Medicine, Baltimore MD 21205 USA; Department of Neuroscience, Haverford College, Haverford, PA 19041 USA; Department of Psychiatry and Behavioral Sciences, Johns Hopkins University School of Medicine, Baltimore MD 21205 USA; Department of Neuroscience, Bryn Mawr College, Bryn Mawr, PA 19010 USA; Kavli Neurodiscovery Institute, Baltimore MD 21218 USA

**Keywords:** Early-life adversity, maternal caregiving, social behavior, amygdala, intercalated cells, limited bedding

## Abstract

Early-life adversity is associated with highly variable developmental outcomes, but identifying sources of this heterogeneity is difficult because variation in outcome is typically accompanied by variation in exposure. Animal models provide experimental leverage by standardizing features of the early environment while allowing variation in developmental experience and outcome to remain observable. Using a graded limited bedding and nesting model spanning normative rearing, moderate restriction, and severe scarcity, we asked what distinguishes social outcomes among animals exposed to the same resource restriction. Severe scarcity increased adverse maternal handling while leaving organized nurturing behaviors intact. Following the P8–P12 rearing manipulation, pups were tested outside the home cage in a P13 social approach assay. Severe scarcity reduced the proportion of pups that approached the dam, but outcomes varied substantially among scarcity litters. Greater typical maternal pup transport predicted greater approach, whereas a greater proportion of rough transport predicted poorer approach. Transport composition did not differ across rearing conditions, but its relationship with social behavior did. Amygdala intercalated-cell recruitment also covaried with social approach under severe scarcity despite limited condition-level differences in regional c-Fos. Plasma corticosterone differed across rearing conditions but did not covary detectably with the principal caregiving, behavioral, or neural measures. Maternal transport and intercalated-cell recruitment were not robustly associated, identifying separable experiential and neural correlates of behavioral variation. These findings show that standardizing an environmental constraint does not produce uniform developmental experience or outcome, and that caregiving experiences preserved across rearing conditions can distinguish social outcomes under severe scarcity.

**Highlights:**

- Severe scarcity increased adverse maternal handling but preserved nurturing care.
- Preserved pup transport predicted later social approach under severe scarcity.
- The transport composition–approach relationship differed by rearing context.
- Intercalated-cell recruitment tracked social approach among scarcity litters.
- Transport and intercalated-cell recruitment were separable behavioral correlates.

## Introduction

Early-life adversity is among the strongest developmental risk factors for anxiety, depression, and stress-related psychiatric disorders (McLaughlin et al., 2019; Teicher & Samson, 2016; Tottenham, 2010), yet developmental outcomes following adversity are highly heterogeneous. Identifying the sources of this heterogeneity is difficult because variation in outcome is typically accompanied by variation in exposure: individuals differ not only in their responses to adversity, but also in the nature, severity, and context of the experiences they encounter. Differences in caregiving, stress-hormone signaling, and the maturation of limbic circuits can each shape developmental outcome (Chen & Baram, 2016; Gee et al., 2013). Animal models provide experimental leverage on this problem by standardizing defined features of the early environment while allowing variation in developmental experience and outcome to remain observable.

Experimental standardization of an environmental constraint does not imply uniformity of the developmental experience that unfolds within it. Experimental effects of adversity are typically characterized by differences between experimental groups, but those comparisons do not capture the variation that emerges within a condition. Animal models make both levels of analysis possible: defined features of the early environment can be controlled while variation in developmental experience and outcome remains measurable. A caregiving behavior that accounts for relatively little behavioral variation under normative conditions, for example, may become more informative under adversity. This makes it possible to distinguish what an experimental manipulation changes on average from what distinguishes outcomes among individuals exposed to the same experimentally defined constraint.

The limited bedding and nesting paradigm provides such an opportunity. Reducing nesting material disrupts maternal caregiving and produces lasting changes in emotional and social behavior (Ivy et al., 2008; Molet et al., 2016; Walker et al., 2017), but scarcity does not uniformly suppress maternal care. Instead, it increases fragmented and disorganized adverse handling (Molet et al., 2016; Rice et al., 2008; Song, McKie, et al., 2026) while leaving organized nurturing behaviors largely intact (Opendak et al., 2019, 2020). Litters exposed to the same resource restriction therefore continue to receive substantial nurturing care that varies from litter to litter, and naturally occurring differences in maternal care can have lasting consequences for offspring phenotype (Champagne et al., 2003). Our previous work further showed that scarcity can preserve nurturing behavior while altering pups’ neural responses to that care through stress hormone-dependent mechanisms (Opendak et al., 2020). Thus, limited bedding standardizes an important dimension of resource scarcity without eliminating potentially meaningful variation in the caregiving environment.

This raises a different question from whether adversity reduces maternal care: *among litters exposed to the same resource restriction, what distinguishes pups that retain caregiver-directed social behavior from those that do not?* Group-level comparisons identify effects associated with adversity, but they do not explain the variation that remains within the adversity condition. Mechanisms associated with preserved behavior also need not simply be the inverse of those associated with impairment. Maternal care is not a unitary developmental input. Distinct caregiving behaviors provide different combinations of nutritional, tactile, vestibular, and social stimulation and can differ substantially in their developmental consequences (Meaney, 2001; Champagne et al., 2003). Moreover, the relationship between a particular caregiving experience and subsequent outcome may depend on the broader rearing context even when the average occurrence of that experience is unchanged. Quantifying the caregiving repertoire therefore permits two related questions to be distinguished: which features of maternal care are altered by resource restriction, and which features distinguish developmental outcomes within that environment.

Understanding this variation also requires identifying neural systems associated with divergent behavioral outcomes. The amygdala is a strong candidate because its contribution to social behavior changes markedly across development (Amaral, 2003), and adversity can alter the maturation of circuitry involved in threat and avoidance (Chen & Baram, 2016; Raineki et al., 2019). Our previous work identified premature recruitment of excitatory basolateral amygdala (BLA) neurons as a mechanism suppressing infant social approach after early-life adversity (Opendak et al., 2021). Here, we ask the complementary question of what neural recruitment distinguishes social outcomes among adversity-exposed animals. Considerably less is known about inhibitory amygdala circuitry during this developmental period. Intercalated cell clusters are GABAergic populations that regulate information flow through the amygdala and contribute to threat regulation and extinction learning in adults (Duvarci & Paré, 2014; Likhtik & Paz, 2015). Because these neurons mature postnatally and gate excitatory amygdala output (Likhtik et al., 2008; Asede et al., 2015), variation in their recruitment may provide a complementary index of developing amygdala function associated with individual differences in social behavior following disrupted caregiving. We also quantified recruitment of the lateral habenula (LHb), which regulates caregiver-directed social behavior in a developmentally changing manner (Cobb-Lewis et al., 2024) and provides a comparison region outside the amygdala for assessing whether brain–behavior relationships were regionally selective.

Here, we used a graded limited-bedding paradigm spanning severe scarcity (100 cc bedding), moderate restriction (500 cc), and normative rearing (2000 cc) to distinguish what resource restriction changed on average from what distinguished outcomes within each rearing condition. We first characterized the maternal caregiving repertoire to identify behaviors altered by resource restriction and those that remained available and variable within rearing conditions. We then asked whether variation in preserved caregiving predicted subsequent caregiver-directed social approach and whether these relationships differed across rearing contexts.

Finally, we assessed amygdala and habenula recruitment following social approach, with particular attention to amygdala intercalated cell clusters as candidate neural correlates of behavioral heterogeneity, and measured plasma corticosterone to determine whether circulating stress hormone levels covaried with the principal caregiving, behavioral, and neural measures.

This design allowed us to examine developmental variation that remained visible within an experimentally standardized environmental constraint.

## Methods

### Animals

All procedures were approved by the Institutional Animal Care and Use Committee and conducted in accordance with NIH guidelines. Long-Evans rats were originally obtained from Envigo and bred in-house. Dams and their litters were housed under standard laboratory conditions (12:12 h light/dark cycle, ad libitum access to food and water) in polypropylene cages (23 L × 20 H × 46 W cm), consistent with standard husbandry practices in developmental stress paradigms (Mroue-Ruiz et al., 2024; Walker et al., 2017). Pups of both sexes were included in approximately equal numbers, and sex was recorded for all pup-level analyses. Litter demographics, dam age, parity, and litter size are summarized in Table S1. Litters were randomly assigned to bedding conditions (see below), and experimenters were blind to condition during pup behavioral scoring and histological quantification, consistent with best-practice recommendations for limited bedding and nesting studies (Ivy et al., 2008; Molet et al., 2016; O’Neill et al., 2025; Walker et al., 2017). Maternal-behavior coders could not be blind to rearing condition, because bedding volume is visible on video; maternal behavior was therefore scored against a predefined ethogram with behavioral definitions fixed *a priori* (see Appendix).

## Bedding manipulation

All pups experienced typical rearing conditions until P8 and again after P12. From P8–P12, litters were assigned to one of three bedding conditions: severe scarcity (100 cc bedding), moderate restriction (500 cc), or normative rearing (2000 cc). The amount and spatial distribution of bedding available under each condition are illustrated in Fig. S1A. This developmental window and manipulation were selected based on prior work showing that restriction of bedding and nesting resources during this period alters maternal caregiving while maintaining pup viability. Bedding volumes were based on established limited bedding and nesting paradigms (Cobb-Lewis et al., 2024; Ivy et al., 2008; Nguyen et al., 2025; Porras et al., 2025; Raineki et al., 2019). Corncob bedding (Inotiv) was used for all conditions. Dams and pups remained undisturbed except for daily video recordings and routine husbandry.

## Behavioral assay

Pup social approach to the dam was assessed on P13 for 30 min using an established pup–dam approach paradigm used to quantify early social behavior following adversity (Raineki et al., 2019; Opendak et al., 2021; see Fig. S1B). The test occurred in a clean polypropylene rat cage (23 L × 20 H × 46 W cm) containing 2000 cc of clean corncob bedding (Inotiv). For testing, an unfamiliar lactating dam of comparable postpartum age to the pup’s biological mother was anesthetized with urethane (2 g/kg) to block milk letdown and positioned laterally to provide pups with access to the nipples. This procedure was consistent with prior pup social approach assays designed to isolate motivational and approach behavior independent of feeding (Opendak et al., 2020; Opendak et al., 2021). Pups were placed individually into the chamber on the side opposite the dam and permitted to interact freely. Approach latencies and zone-specific contact durations were quantified from video by trained coders using EthoVision XT (Noldus), consistent with prior work using this assay (Opendak et al., 2021).

## Immunohistochemistry and c-Fos quantification

Pups were placed into a quiet, covered container on a heating pad and perfused 60 min after the social approach test, a time point chosen to capture immediate early gene expression following behavioral engagement (Herrera & Robertson, 1996; Kovács, 1998). Following deep isoflurane anesthesia, animals were transcardially perfused with cold PBS followed by 4% paraformaldehyde (PFA). Brains were post-fixed in 4% PFA for 24 h, cryoprotected in 30% sucrose, and coronally sectioned at 40 μm using standard protocols for c-Fos immunohistochemistry in developing rodent brain (Raineki et al., 2019; Cobb-Lewis et al., 2024). Sections were washed in PBS, quenched in 3% H₂O₂, blocked in 3% normal horse serum, and incubated in primary antibody (rabbit anti-c-Fos, 1:5000; Invitrogen MA5-15055) for 44–48 h at 4°C, consistent with established c-Fos labeling protocols (Barth et al., 2004; Kovács, 1998). After PBS washes, sections were incubated in biotinylated secondary antibody (1:500; Vector BA-1100-1), amplified using the ABC-HRP kit (Vector PK-4000), and developed using nickel-enhanced DAB (Sigma D12384), as commonly used for high-contrast c-Fos visualization (Barth et al., 2004; Song, McKie et al., 2026). Sections were counterstained with methyl green, dehydrated, cleared, and coverslipped with DPX.

Fos-positive cells were quantified bilaterally in the basolateral amygdala (BLA), dorsal intercalated cell clusters (dITC), ventral intercalated cell clusters (vITC), and lateral habenula (LHb), based on established anatomical boundaries and nomenclature for these regions (Cobb-Lewis et al., 2024; Duvarci & Paré, 2014; Likhtik et al., 2008; Opendak et al., 2021), using ImageJ (NIH; Schindelin et al., 2012; Song, McKie et al., 2026). Regions of interest were delineated anatomically, thresholded particles of 10–150 μm² were counted, and counts were summed across six sections and normalized by sampled tissue area to obtain regional c-Fos densities (cells/mm²). Combined left- and right-hemisphere values were used for all analyses. Threshold settings and the blinded adjustment rule are given in the Supplementary Methods.

## Corticosterone assay

Trunk blood was collected from P12 littermates, and plasma corticosterone was quantified by ELISA (four litters per rearing condition; Enzo Life Sciences) according to the manufacturer’s protocol, except as noted below. Samples were assayed in duplicate at a 1:20 dilution to ensure that all values fell within the linear range. Corticosterone was analyzed at the litter level.

## Maternal behavior

Maternal behavior was recorded for 1 h daily during the manipulation window (P8–P12) and scored from video in BORIS (Friard & Gamba, 2016) using a predefined 23-behavior ethogram (22 analyzed; see Appendix). Each behavior was summarized per recording in two ways: as the proportion of 10-min bins in which it occurred (scan proportion) and as the percentage of session time it occupied (percent duration). Percent duration was the primary measure for sustained states such as nest occupancy and nursing; scan proportion was primary for brief discrete events such as pup transport, whose durations fall near the 0.1% storage resolution of the duration measure. Recordings were obtained and analyzed across five days in 21 litters and four days in three litters. Litter means were computed over the sessions available for each measure rather than over a nominal session count. Rough transport share was computed per litter as rough transport divided by the sum of rough and typical transport, separately for each measure, and was undefined for litters with no scored transport. Condition and day effects on each of the 22 behaviors were modeled with generalized estimating equations clustered by litter, with false-discovery-rate correction of the condition × day term across behaviors (Table S2). An exploratory analysis of the sequential unpredictability of maternal behavior (transition entropy) was conducted on the subset of sessions with event-level records. Both procedures are described in the Supplementary Methods.

## Pup social approach

Approach latency to the dam was quantified per pup as the minimum latency across the three reachable contact zones and treated as right-censored at the maximum session duration (1800 s) for pups that did not approach. Because the caregiving predictors of interest (pup transport and stepping) were litter-level variables, approach was summarized per litter using two measures: the proportion of pups in each litter that approached and the restricted mean survival time (RMST) of that litter’s Kaplan–Meier curve over the 1800-s test. RMST is the area under the curve for not yet having approached, expressed in seconds, and is interpretable as the average time a pup in that litter spent without having approached the dam (Irwin, 1949; Royston & Parmar, 2011). It ranges from 0 s, when every pup approaches immediately, to 1800 s, when no pup approaches; lower values therefore indicate faster and more complete approach. Unlike mean or median latency, RMST remains defined when some pups never approach and incorporates both the timing and occurrence of approach.

Differences in approach timing between conditions were assessed with Kaplan–Meier estimates and log-rank tests (Kaplan & Meier, 1958); the proportional-hazards assumption was not rejected (Supplementary Methods). Analyses were conducted in the full behavioral cohort and, separately, in the subset of pups processed for c-Fos, and condition contrasts are labeled accordingly. Because c-Fos was measured in two pups per litter, litter-level neural analyses used the geometric-mean approach latency of those two pups, whereas caregiving analyses used RMST across all tested pups in the litter; within severe scarcity, the two summaries were rank-identical for the imaged pups. The degree to which approach was structured by litter was quantified with intraclass correlation coefficients, pup-level Cox models with a caregiving predictor × severe-scarcity interaction were fitted as a complement to the litter-level correlations, and zone-occupancy phenotypes were derived descriptively by clustering; these procedures are described in the Supplementary Methods.

## c-Fos analysis

Fos-positive cells were quantified bilaterally and expressed as density (counts per sampled area, summed across sections). Regional sample sizes were 48 pups for BLA, dITC, and LHb and 44 pups for vITC, for which clusters were absent from recovered sections or could not be delineated reliably in four pups spanning all three rearing conditions. Analyses used two processed pups from each litter, selected to be behaviorally representative: where possible, one male and one female whose latency to the ventrum fell within approximately one standard deviation of the litter mean, without reference to c-Fos data.

Condition differences were tested separately for each region using Kruskal–Wallis tests; these analyses characterized condition-level regional recruitment and were not part of the within-severe-scarcity brain–behavior family. Within severe scarcity, regional c-Fos was correlated (Spearman) at the litter level with censored minimum approach latency and with two a priori composites of approach speed and contact time (Supplementary Methods), using litter-resampled confidence intervals and exact permutation p-values, and results for all regions are reported. Condition dependence of the c-Fos associations, joint models of transport and dITC c-Fos, and within-litter sibling contrasts are described in the Supplementary Methods. Two directional tests were specified in advance. BLA c-Fos was predicted to be higher under severe scarcity than normative rearing based on prior work in this model (Opendak et al., 2021). Because intercalated cells gate excitatory amygdala output, the association between adverse maternal handling (stepping on pups) and dITC c-Fos was predicted to be negative; this association was additionally estimated with rearing condition partialled out. The severe-scarcity ICC contrast described above was also evaluated directionally. All other comparisons were two-tailed.

Because the LHb has been implicated in the development of caregiver-directed social behavior, the seven organized nurturing behaviors quantified here were tested against LHb c-Fos as a defined multiple-comparison family with Benjamini–Hochberg correction across the seven tests (Supplementary Methods). A wider screen of all maternal behaviors against all four regions was exploratory and is reported in Supplementary Analysis 7.

## Statistical analysis

Analyses were performed in Python (statsmodels, scipy, lifelines) and R. Litter was the unit of inference for litter-level variables, including maternal behavior and corticosterone. Pup-level measures, including individual approach latency and c-Fos, were analyzed at the pup level where appropriate, with within-litter dependence accounted for through clustering, litter-level permutation, or litter-level summaries as specified above; litter-level predictors were not regressed on pup-level observations without accounting for clustering (Lazic, 2010; Lazic & Essioux, 2013). Because several analyses included eight litters per condition, inference did not rely on asymptotic p-values: litter-level correlations within a rearing condition were tested by exact permutation over all relabelings of that condition’s litters (8! = 40,320; 7! = 5,040 where one litter recorded no transport and its rough share was therefore undefined), pup-level statistics involving a litter-level predictor were tested by permuting whole litters, and leave-one-litter-out analyses assessed the sensitivity of the principal caregiving association to individual litters. Cluster-robust and bootstrap procedures, and the design effects of pup-level analyses, are detailed in the Supplementary Methods. All tests were two-tailed except the directional BLA, stepping–dITC, and ICC contrasts described above. The rough-share moderation, the condition effect on approach, and condition differences in litter clustering were treated as exploratory or descriptive and are identified as such in the Results.

## Results

### Severe early-life scarcity is associated with reduced infant social approach

We first asked whether graded resource restriction altered caregiver-directed social approach during infancy (Fig. 1B). In the balanced neural cohort, pups exposed to severe scarcity showed reduced approach relative to normative-reared pups (log-rank χ² = 4.32, *p =* 0.038), whereas moderate restriction did not differ from normative rearing (*p =* 0.57). Analysis of the larger behavioral cohort showed a similar descriptive pattern (Supplementary Analysis 1). The severe-scarcity phenotype was driven primarily by an increased proportion of pups that did not approach within the 30-min test rather than by differences in approach latency among pups that approached. Rearing conditions did not differ detectably in body weight, litter size, dam demographics, or spatial occupancy during the test (Table S1; Fig. S1; Supplementary Analysis 1).

**Figure 1.**
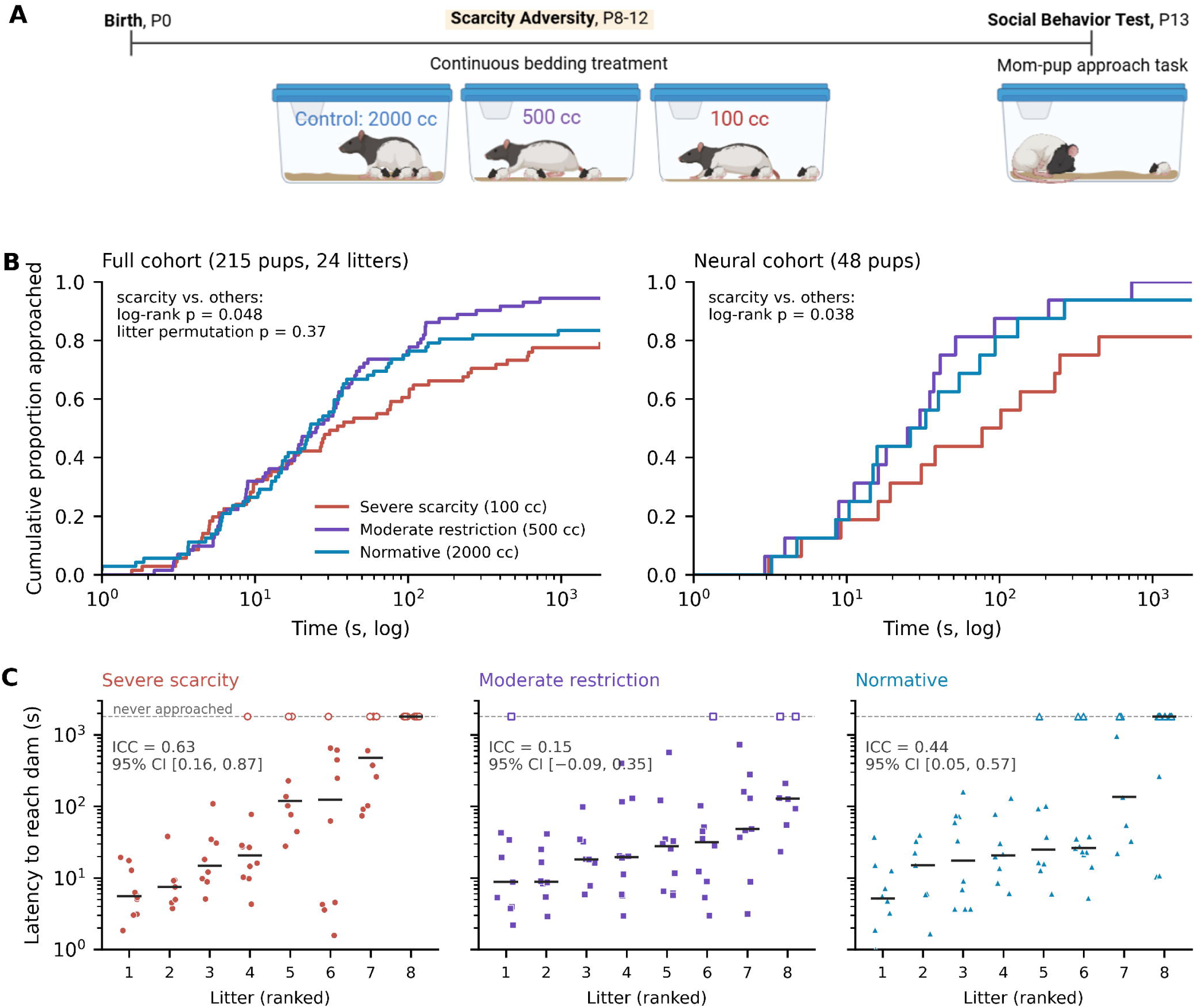
Severe scarcity reduces the likelihood that pups reach the dam and is where approach is most strongly structured by litter. (A) Experimental timeline. Litters were reared under one of three bedding conditions, normative (2000 cc), moderate restriction (500 cc), or severe scarcity (100 cc), continuously from postnatal day (P) 8 to P12, and pup social approach to the anesthetized dam was assessed on P13 (apparatus and scored zones in Fig. S1). (B) Cumulative incidence of approach (Kaplan–Meier estimates, log time axis) for the full behavioral cohort (215 pups, 24 litters) and the neural cohort (48 pups, two per litter), with pups not approaching within the 1800 s test treated as right-censored. Because bedding condition was assigned to litters, inference is drawn from the neural cohort, in which every litter contributes exactly two pups and clustering across the pooled cohort is slight (design effect 1.47 on binary approach): approach was slower under severe scarcity than under the two milder conditions combined (log-rank *p =* 0.038), whereas the three-condition omnibus test was not significant (*p =* 0.107). The full behavioral cohort is shown descriptively, since its pups are strongly clustered (binary approach ICC = 0.39, mean 9.0 pups per litter, design effect 4.1); its nominal log-rank values (*p =* 0.048 for severe scarcity versus the other conditions; *p =* 0.071 omnibus) are reported alongside the same statistics evaluated by permuting rearing condition across whole litters (*p =* 0.38 and *p =* 0.60; 20,000 permutations). (C) Approach latency for every pup in the full cohort, grouped by litter within each rearing condition. Litters are ranked by median latency; horizontal bars are litter medians; open symbols are pups that did not approach, plotted at the 1800 s ceiling. The value in each panel is the proportion of variance in log10 approach latency attributable to litter (one-way ICC(1); non-approaching pups entered at 1800 s) with its 95% confidence interval from bootstrap resampling of litters. Litter clustering was highest under severe scarcity (0.63; normative rearing 0.44; moderate restriction 0.15), and the same ordering held in the neural cohort (0.72, 0.14 and 0.53); severe scarcity versus the two milder conditions, litter-permutation *p =* 0.087 (full cohort) and 0.083 (neural cohort).

The proportion of variation in social approach attributable to litter was descriptively highest under severe scarcity (Fig. 1C). Intraclass correlation coefficients for pup approach latency (log-transformed, censored at 1800 s; Supplementary Methods) were highest under severe scarcity (ICC = 0.63, 95% CI 0.16 to 0.87), intermediate under normative rearing (ICC = 0.44, 95% CI 0.05 to 0.57), and lowest under moderate restriction (ICC = 0.15, 95% CI −0.09 to 0.35); the contrast between severe scarcity and the two milder conditions did not reach significance (ΔICC = 0.31, litter-permutation *p =* 0.087). Severe scarcity again showed the highest litter clustering in the neural cohort (ICC = 0.72, 0.53, and 0.14 for severe scarcity, moderate restriction, and normative rearing; severe scarcity versus the other two conditions, *p =* 0.083), and the severe-scarcity estimate in the full cohort remained the highest of the three when any single litter was omitted (range 0.48–0.77). Thus, despite standardized bedding availability within the severe-scarcity condition, social approach was strongly structured by litter. This pattern motivated subsequent analyses asking whether variation in maternal caregiving distinguished behavioral outcomes among litters exposed to the same resource restriction.

## Scarcity increases adverse maternal handling while preserving organized nurturing care

We next characterized maternal behavior across the graded resource-restriction conditions (Fig. 2; Table S2). Severe scarcity selectively increased adverse maternal handling, with stepping on pups showing a significant condition × day interaction in the logistic GEE model (*p =* 0.0004, *q =* 0.007). Across sessions, the proportion of scans containing stepping increased from 0.26 under normative rearing to 0.57 under moderate restriction and 0.73 under severe scarcity, and stepping increased with the severity of resource restriction (Spearman ρ = 0.60, *p =* 0.002).

**Figure 2.**
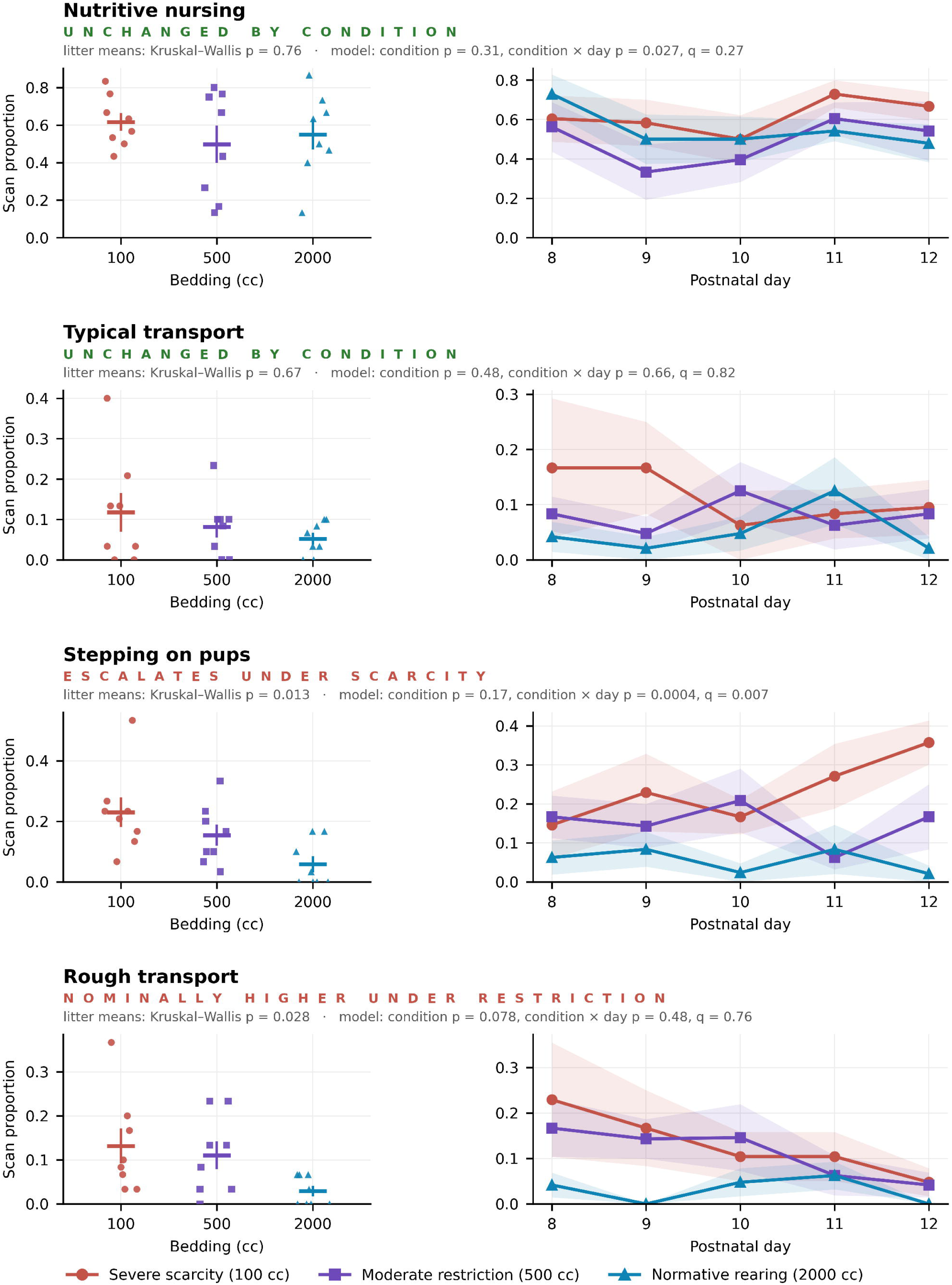
Early-life scarcity intensifies adverse maternal handling while leaving organized nurturing care unchanged. Maternal behavior was scored from daily one-hour recordings across the manipulation window (postnatal day 8–12) using a 23-behavior ethogram (see Appendix). For each behavior, the left panel shows litter means by rearing condition (points, individual litters; horizontal bars, condition mean ± SEM) and the right panel the day-resolved trajectory (line, condition mean; band, ± SEM). All four behaviors are shown on the scan-proportion measure. (Top two rows) Nutritive nursing and typical pup transport did not differ across rearing conditions, and neither showed a day-resolved divergence surviving correction. (Bottom two rows) Maternal stepping on pups escalated across the exposure window specifically under severe scarcity (condition × day *p =* 0.0004, *q =* 0.007), the only maternal effect surviving false-discovery-rate correction. Rough pup transport was elevated under restriction on litter means (Kruskal–Wallis *p =* 0.028) but produced no day-resolved divergence in the model of Table S2 (condition × day *p =* 0.48, *q =* 0.76). Statistics annotated on each row are the Kruskal–Wallis test on litter means and the condition and condition × day terms of that behavior’s model in Table S2. Vertical axes are scaled separately for each behavior and are not comparable between rows.

In contrast, organized nurturing behaviors remained largely intact across conditions. None of the organized nurturing measures differed significantly across rearing conditions (all p ≥ 0.19; Fig. 2; Fig. S2C–D; Supplementary Analyses 2–3; Supplementary Methods). Thus, the principal detectable change in maternal behavior was an increase in adverse handling rather than a generalized reduction in organized nurturing care.

## Variation in preserved maternal transport predicts subsequent social approach under severe scarcity

Because organized nurturing behaviors remained available under severe scarcity, we next asked whether variation in this preserved care distinguished subsequent social outcomes among litters exposed to the same resource restriction. Within severe scarcity, litters whose dams engaged in more typical pup transport subsequently showed greater caregiver-directed social approach (scan proportion: ρ = −0.75, exact *p =* 0.040; transport bouts: ρ = −0.83, exact *p =* 0.017; Fig. 3A). The association was stable in direction and magnitude across leave-one-litter-out analyses (ρ = −0.67 to −0.80), although removal of litter 26-60 reduced the association to *p =* 0.11 (ρ = −0.67; Fig. S4; Supplementary Analysis 4).

**Figure 3.**
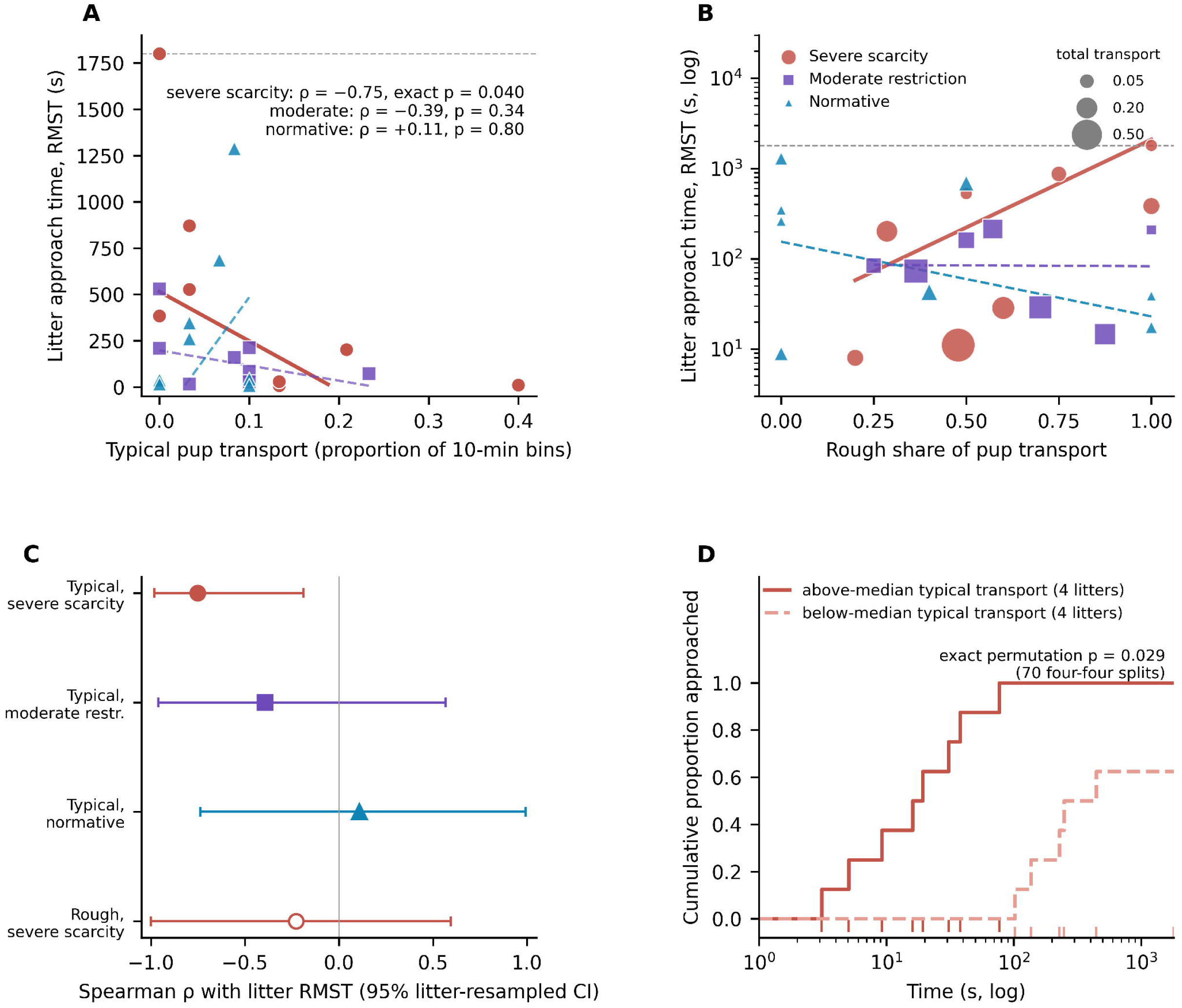
Variation in typical pup transport, and in the composition of transport, predicts infant social approach under severe scarcity. (A) Litter-level typical transport against the approach outcome, by rearing condition. The approach outcome is the restricted mean survival time (RMST) of each litter’s Kaplan–Meier curve over the 1800 s test, so lower values indicate faster and more complete approach; lines are Theil–Sen fits within each condition, truncated where they would predict a negative RMST. Within severe scarcity (solid line), Spearman ρ = −0.75, exact permutation *p =* 0.040 over all 40,320 relabelings of the eight litters; no comparable relationship was present under moderate restriction (ρ = −0.39, *p =* 0.34) or normative rearing (ρ = +0.11, *p =* 0.80; dashed lines). (B) The composition of transport, expressed as the rough share of all scored transport, against the same approach outcome for all litters with any scored transport (23 of 24; one litter recorded no transport, so the share is undefined), by rearing condition. Lines are Theil–Sen fits on log RMST within each condition. Marker area is the total transport scored, the denominator of the share, so litters at a share of zero or one rest on the fewest observations and the moderate-restriction and normative slopes are correspondingly coarse. Within severe scarcity the share was associated with slower approach (Spearman ρ = +0.75, exact permutation *p =* 0.038), whereas it was unrelated to approach under moderate restriction (ρ = −0.04) and normative rearing (ρ = −0.32) and across all litters (ρ = +0.06, *p =* 0.77); the severe-scarcity × rough-share interaction was significant (*p =* 0.007; Supplementary Analysis 4). (C) Spearman ρ with litter-resampled 95% confidence intervals for typical transport within each rearing condition, and for rough pup transport within severe scarcity. Only typical transport under severe scarcity excludes zero, indicating that the relationship is concentrated under scarcity and specific to typical rather than rough handling. (D) Kaplan–Meier approach curves in the neural cohort (two pups per litter, four litters per group), with litters split at the median transport level; rug marks show individual pups. Two pups per litter keep the design effect near 1.5, and the split is tested by exact permutation over all 70 possible four-four divisions of the eight litters (*p =* 0.029).

Transport organization also provided information about subsequent behavior. Within severe scarcity, a greater proportion of transport events classified as rough was associated with poorer subsequent approach (ρ = 0.75, *p =* 0.038), as was the proportion of transport duration that was rough (ρ = 0.83, *p =* 0.030; Fig. 3B). Absolute rough transport, however, was not associated with approach, suggesting that the composition of maternal transport rather than the amount of rough transport alone carried the behavioral association.

The relationship between rough transport share and social approach differed across rearing conditions (interaction *p =* 0.007), despite no overall condition difference in rough transport share (medians 0.55, 0.57, and 0.20; Kruskal–Wallis *p =* 0.33). This interaction remained significant across alternate specifications and leave-one-litter-out analyses (all *p* ≤ 0.040). A pup-level Cox model (Supplementary Methods) provided convergent evidence for condition dependence (interaction log-HR = −1.20, 95% CI −2.52 to −0.21, *p =* 0.016); within severe scarcity, each SD increase in rough transport share was associated with a lower rate of approach (HR = 0.42, 95% CI 0.14 to 0.81). In contrast, the formal interaction for typical transport did not reach significance (*p =* 0.20).

Typical transport and rough transport share were themselves strongly related within severe scarcity (ρ = −0.82), and neither showed an independent association with approach when entered together. Because the rough-share measure was exploratory and in some litters was based on relatively few transport events, we retain typical transport as the prespecified primary measure and transport composition as convergent, exploratory evidence that the relationship between maternal transport and subsequent social behavior depends on rearing context.

## Amygdala intercalated-cell recruitment covaries with litter-level variation in social approach

We next asked whether behavioral variation was accompanied by variation in neural recruitment following the social approach test. A prespecified directional comparison detected greater BLA recruitment under severe scarcity than normative rearing (one-tailed *p =* 0.047). Overall c-Fos expression did not differ significantly across rearing conditions in ventral intercalated cells (vITC; *p =* 0.074), dorsal intercalated cells (dITC; *p =* 0.86), or LHb (*p =* 0.17; Fig. 4B–E).

**Figure 4.**
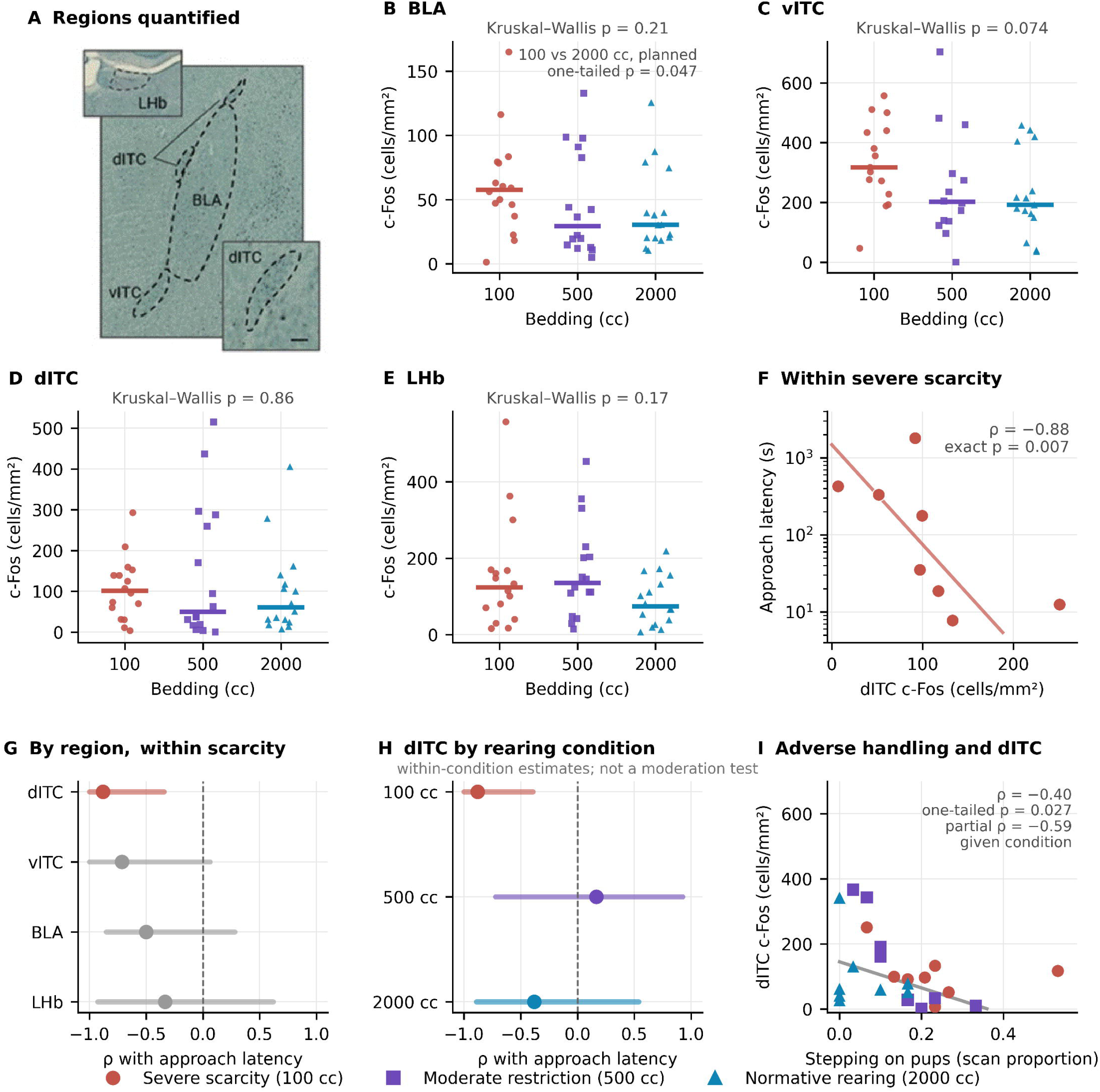
Intercalated cell activity tracks infant approach under severe scarcity. Estimates in (B–E) are at the pup level, as specified for c-Fos in the Methods; those in (F–I) are at the litter level. (A) Representative section showing the regions quantified: basolateral amygdala (BLA), ventral and dorsal intercalated cell clusters (vITC, dITC), and lateral habenula (LHb); scale bar, 100 µm. (B–E) c-Fos density (cells/mm²) by rearing condition; points are individual pups, bars show condition medians. No region showed an omnibus difference between rearing conditions (Kruskal–Wallis, all p ≥ 0.074), indicating that scarcity did not produce generalized changes in circuit activity. In the single contrast specified in advance, BLA c-Fos was higher under severe scarcity than under normative rearing (one-tailed *p =* 0.047; panel B). (F) dITC c-Fos against approach latency within severe scarcity (ρ = −0.88, exact permutation *p =* 0.007; each point is a litter). (G) Spearman correlations of regional c-Fos with approach latency under severe scarcity (points, ρ; bars, 95% confidence interval from litter resampling); dITC was the only region whose interval excluded zero, although the amygdala subregions are collinear and no region survives correction across the full set of region-by-outcome tests. (H) The dITC–approach association estimated separately within each rearing condition. These are within-condition estimates and not a test of moderation: the formal condition × dITC interaction does not reach significance on log-transformed densities and is sensitive to model specification (Results). (I) Maternal stepping on pups against dITC c-Fos across all 24 litters (ρ = −0.40, one-tailed permutation *p =* 0.027 for the directional prediction stated in the Results; partial ρ = −0.59 adjusting for rearing condition). No comparable relationship was present in vITC (ρ = −0.01), BLA (ρ = −0.28), or LHb (ρ = −0.09).

Despite limited condition-level differences in regional c-Fos, dITC recruitment covaried strongly with social approach among severe-scarcity litters (ρ = −0.88, 95% CI −1.00 to −0.39, exact *p =* 0.007; Fig. 4F–G). The same relationship was evident using the approach-speed (*p =* 0.022) and contact-time (*p =* 0.037) composites (Supplementary Methods). The dITC association remained strong after accounting for BLA recruitment (partial ρ = −0.84), whereas the corresponding association between BLA recruitment and approach was attenuated after accounting for dITC recruitment (partial ρ = 0.06). A similar analysis comparing vITC and BLA yielded partial correlations of −.59 and −.01, respectively (Supplementary Analysis 5). These analyses identify intercalated-cell recruitment as a neural correlate of behavioral variation but do not establish anatomical specificity.

Maternal stepping on pups was also associated with lower dITC recruitment across litters (ρ = −0.40, one-tailed permutation *p =* 0.027), and this association remained after accounting for rearing condition (partial ρ = −0.59; Fig. 4I). Stepping itself, however, was not associated with social approach (ρ = 0.07, *p =* 0.73).

## Maternal transport and intercalated-cell recruitment are separable correlates of social behavior

Because both typical maternal transport and dITC recruitment covaried with social approach, we asked whether they were themselves related. Typical transport was not detectably associated with dITC recruitment across all litters (ρ = −0.08, *p =* 0.72) or within severe scarcity (ρ = 0.61, exact *p =* 0.12). Joint models were limited by sample size and did not provide evidence that dITC recruitment accounted for the transport–approach association (Supplementary Analysis 6).

We additionally examined whether differences in dITC recruitment between littermates tracked differences in their social approach. Within-litter differences in dITC recruitment were not systematically associated with within-litter differences in approach (Wilcoxon *p =* 0.45; Supplementary Methods); only three of seven severe-scarcity pairs showed the predicted ordering, the eighth pair being uninformative because neither imaged pup approached (Supplementary Analysis 5). Thus, the dITC–approach association was evident between litters but was not detected within littermate pairs. The present data therefore do not support a transport → dITC → approach pathway. Typical maternal transport and intercalated-cell recruitment are instead treated as separable correlates of social behavior whose relationship to one another cannot be resolved by the present design. Because c-Fos was measured after the approach test, neural recruitment additionally reflects circuit engagement associated with behavior rather than a temporally antecedent mediator.

## Maternal milk ejection covaries with LHb recruitment across rearing conditions

A broader analysis of organized nurturing behaviors identified an association between maternal milk ejection and LHb recruitment. Across rearing conditions, greater time spent in milk ejection was associated with lower LHb c-Fos (standardized β = −0.54; Spearman ρ = −0.56, *p =* 0.005, *q =* 0.032; exact permutation *p =* 0.005; Fig. 5; family defined in the Supplementary Methods). None of the other organized nurturing behaviors tested showed a detectable relationship with LHb recruitment (all p ≥ 0.22).

**Figure 5.**
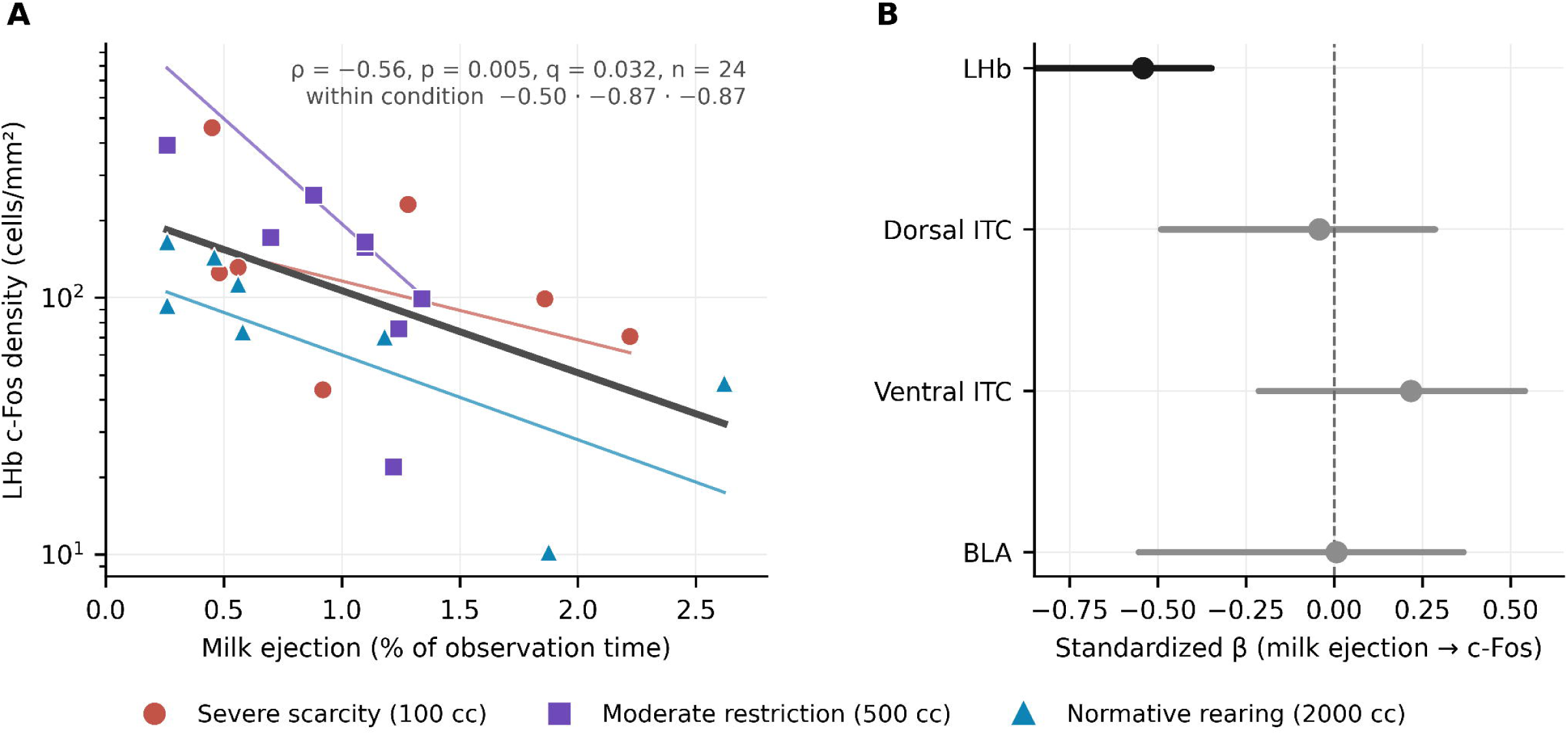
A caregiving–habenula relationship that does not depend on rearing condition. All estimates are at the litter level; c-Fos densities were log-transformed for analysis. (A) Litter mean milk ejection, as a percentage of observation time across the P8–12 recording window, against LHb c-Fos density measured after the P13 approach test (24 litters; log axis). Litters in which milk ejection was more prolonged showed lower habenular recruitment (Spearman ρ = −0.56, *p =* 0.005; Pearson r = −0.54, *p =* 0.006; permutation *p =* 0.005 over 200,000 relabelings; *q =* 0.032 across the declared family of seven organized nurturing behaviors tested against LHb c-Fos). The association was present on percent duration and not on the scan-proportion measure of the same behavior (ρ = −0.17, *p =* 0.43). The relationship was present within each rearing condition (thin lines; ρ = −0.50 under severe scarcity, −0.87 under moderate restriction, −0.87 under normative rearing), and neither variable differed between conditions (milk ejection Kruskal-Wallis *p =* 0.71; LHb c-Fos *p =* 0.24). Adjusting for rearing condition strengthened rather than attenuated the association (standardized β = −0.57, *p =* 0.004), and it is likewise not attributable to pup sex (Results; Fig. S4E–F). Thick line, fit across all litters. (B) Standardized coefficients for milk ejection predicting c-Fos in each region quantified (points, β; bars, 95% confidence interval from litter resampling). Only the LHb interval excluded zero (β = −0.54, CI [−0.88, −0.35]); dorsal ITC (β = −0.04, CI [−0.49, +0.29]), ventral ITC (β = +0.22, CI [−0.21, +0.54]) and BLA (β = +0.01, CI [−0.55, +0.37]) showed no relationship. LHb c-Fos was not significantly associated with approach in these litters (against geometric-mean latency of the imaged pups, ρ = −0.18, *p =* 0.40; against litter RMST across all tested pups, ρ = −0.35, *p =* 0.09), so this relationship is separate from the scarcity-dependent intercalated-cell associations of Figure 4 rather than a further step in the same pathway.

This association depended on how milk ejection was quantified: percent duration was associated with LHb recruitment, whereas scan proportion was not (ρ = −0.17, *p =* 0.43), although the two measures were themselves correlated (ρ = 0.61). Neither milk ejection nor LHb c-Fos differed significantly across rearing conditions, and LHb recruitment was not significantly associated with social approach (geometric-mean latency ρ = −0.18, *p =* 0.40; Supplementary Analysis 7). The milk ejection–LHb relationship therefore appears distinct from the scarcity-associated transport–approach and dITC–approach relationships.

## Corticosterone does not covary detectably with the principal caregiving, behavioral, or neural measures

Finally, we asked whether circulating corticosterone covaried with the principal caregiving, behavioral, or neural measures. Plasma corticosterone measured in P12 littermates differed across rearing conditions (Kruskal–Wallis *p =* 0.044), although the pattern was nonmonotonic with resource restriction. Corticosterone did not correlate detectably with typical maternal transport, social approach, or dITC recruitment (Supplementary Analysis 8). Because corticosterone was measured in separate littermates and the sample was limited, these analyses do not establish independence from stress-hormone signaling. Rather, variation in circulating corticosterone measured at P12 did not recapitulate the principal caregiving, behavioral, or neural relationships observed here.

## Discussion

The present study asked what distinguishes developmental outcomes among litters exposed to the same experimentally imposed resource restriction. Although bedding availability was standardized within each condition, the caregiving experiences that unfolded under that constraint were not. Severe scarcity increased adverse maternal handling, particularly stepping on pups, while leaving organized nurturing behaviors largely intact. Yet the caregiving features most clearly altered by scarcity were not those that distinguished subsequent social outcomes. Typical maternal pup transport predicted caregiver-directed social approach among severe-scarcity litters, and the relationship between transport composition and behavior differed across rearing contexts. Social approach also varied substantially among litters under severe scarcity, and amygdala intercalated-cell recruitment covaried with this behavioral variation despite limited condition-level differences in regional c-Fos. These findings show that standardizing resource availability does not produce uniform developmental experience or outcome.

Resource restriction reliably increases fragmented, unpredictable, and adverse maternal behavior without necessarily producing a corresponding reduction in nurturing care (Ivy et al., 2008; Rice et al., 2008; Molet et al., 2016; Walker et al., 2017; Opendak et al., 2019, 2020; Song, McKie, et al., 2026). We observed the same dissociation: severe scarcity increased stepping on pups but did not broadly reduce organized nurturing behavior. More importantly, the behavior most clearly altered by the manipulation was not the behavior most strongly associated with subsequent social approach. Stepping increased with resource restriction but did not predict approach, whereas typical transport did not differ across rearing conditions yet predicted approach under severe scarcity. Thus, the features of caregiving that characterize an adverse environment at the group level need not be those that distinguish outcomes within it.

Maternal care comprises distinct nutritional, tactile, vestibular, and social inputs with different developmental consequences (Meaney, 2001; Champagne et al., 2003). Pup transport was particularly informative because it remained available across rearing conditions while varying considerably among litters. Under severe scarcity, greater typical transport was associated with greater subsequent caregiver-directed approach, whereas a greater proportion of rough transport was associated with poorer approach. Transport composition itself did not differ across rearing conditions, but its relationship with social behavior did. Scarcity therefore changed the context in which variation in maternal transport was related to subsequent behavior rather than simply changing transport itself. This extends our previous finding that scarcity rearing altered pups’ neural responses to nurturing maternal care even when that care remained available (Opendak et al., 2020). In both studies, the broader rearing environment changed how preserved caregiving experience was related to the developing pup.

The litter-level structure of social approach reinforces this point. A substantial proportion of variation in approach was attributable to litter under severe scarcity, and this pattern was stable to removal of individual litters. Litter membership encompasses maternal behavior, sibling interactions, biological factors, and other features of the shared environment (Lazic, 2010; Lazic & Essioux, 2013), so the ICC does not identify the source of this variation. It does show that pups exposed to the same experimentally standardized resource restriction did not converge on a uniform behavioral phenotype. Maternal transport identifies one measurable feature that varied among these litters and predicted subsequent behavior. Experimental control over the environmental constraint therefore exposed rather than eliminated variation in the developmental experiences occurring within it.

Our previous work approached this phenotype from a different direction. Early-life adversity increased the proportion of pups that failed to approach the caregiver, and suppressing BLA activity restored approach in adversity-exposed pups (Opendak et al., 2021). That work identified a circuit mechanism contributing to adversity-induced social avoidance, but did not ask why some pups continued to approach despite adversity. Here, we asked that complementary question: *what distinguishes pups that retain social approach under adverse rearing?* Mechanisms supporting preserved behavior need not simply be the inverse of those producing impairment. Consistent with this distinction, BLA recruitment was not the strongest neural correlate of variation within severe scarcity; intercalated-cell recruitment was. At the level of caregiving experience, preserved maternal transport distinguished behavioral outcomes. Focusing on within-condition variation therefore revealed experiential and neural features associated with preserved social behavior that are obscured by the mean effect of adversity alone.

Intercalated-cell c-Fos covaried strongly with social approach among severe-scarcity litters despite limited overall effects of rearing condition on regional c-Fos. dITC showed the strongest regional association, and its relationship with behavior remained after accounting for BLA recruitment, whereas the BLA association was attenuated after accounting for dITC. Intercalated cells regulate information flow through the amygdala and provide inhibitory control over excitatory amygdala output (Likhtik et al., 2008; Asede et al., 2015). In adults, these GABAergic clusters contribute to threat regulation and extinction learning (Duvarci & Paré, 2014; Likhtik & Paz, 2015), but their role during early development is considerably less understood. Their association with caregiver approach is particularly interesting during a period when amygdala contributions to caregiver-directed behavior are changing and adversity can prematurely recruit BLA circuitry associated with avoidance (Raineki et al., 2019; Opendak et al., 2021). Intercalated cells therefore emerge as a candidate component of developing amygdala circuitry associated with variation in infant social behavior following adversity.

The littermate analysis further localized this relationship. Differences in dITC recruitment between siblings did not track differences in social approach or time in contact with the dam (Supplementary Analysis 5); instead, the dITC–approach association was evident between litters. This parallels the litter-level structure of the behavioral phenotype and raises the possibility that dITC recruitment reflects developmental influences operating at the level of the shared litter environment. At the same time, maternal transport and dITC recruitment were not robustly associated despite both covarying with social approach under severe scarcity. Maternal stepping showed yet another pattern, predicting lower dITC recruitment but not social approach. Maternal transport and intercalated-cell recruitment therefore identify separable experiential and neural correlates of behavioral variation rather than successive steps in a single caregiving–neural pathway.

Additional measures identified relationships distinct from this principal phenotype. Greater maternal milk ejection was associated with lower LHb c-Fos across rearing conditions, despite no condition differences in either measure and no relationship between LHb recruitment and social approach. The contribution of the LHb to caregiver-directed social behavior changes markedly across early development (Cobb-Lewis et al., 2024), making this caregiving–LHb association an interesting target for future study, but its presence across rearing conditions distinguishes it from the scarcity-associated transport and intercalated-cell relationships. Plasma corticosterone likewise differed across rearing conditions but did not covary detectably with maternal transport, intercalated-cell recruitment, or social approach. Previous work showed that stress hormones can alter pups’ neural responses to maternal care under scarcity (Opendak et al., 2020); the present results indicate that a single P12 circulating corticosterone measure did not capture the variation associated with the behavioral phenotype.

Several features define the scope of these findings. The principal caregiving analyses included eight litters per condition, so we used exact litter-level permutation, cluster-aware inference, and leave-one-litter-out analyses to assess their stability. Differences in predictor range limited formal tests of moderation for typical transport, whereas transport composition provided stronger evidence that the relationship with behavior differed across rearing contexts. The ICC identifies shared litter-level structure without distinguishing maternal, sibling, genetic, or other biological and environmental contributions. These considerations constrain mechanistic interpretation but do not alter the central observation that substantial experiential, behavioral, and neural variation emerged within an experimentally standardized resource environment.

Developmental outcomes following early-life adversity are heterogeneous in both humans and experimental models (Brenhouse & Bath, 2019; Goodwill et al., 2019; Gee, 2021; McLaughlin et al., 2021). Human studies face the additional challenge that adverse experiences vary in their nature, severity, and context, making it difficult to separate heterogeneity in exposure from heterogeneity in outcome. Experimental models provide complementary leverage by constraining defined features of the early environment while measuring the variation that emerges within them. Here, standardized resource restriction did not produce uniform developmental experience or behavioral outcome. Severe scarcity increased adverse maternal handling, but caregiving experiences preserved across rearing conditions distinguished social outcomes under severe scarcity, and variation in amygdala intercalated-cell recruitment tracked the same behavioral phenotype. The features that define an adverse environment at the group level are therefore not necessarily those that distinguish outcomes within it. Understanding developmental heterogeneity after adversity requires attention to the experiences that continue to vary within an adverse environment, not only those that adversity changes on average

## Supporting information

Source data and statistics

## CRediT authorship contribution statement

Allison Su: Conceptualization, Investigation, Formal analysis, Writing – review & editing. Elaina Regier: Investigation, Writing – review & editing. Chloe Lim: Investigation, Writing – review & editing. Isaac McKie: Investigation, Writing – review & editing. Ben Perez-Flesler: Investigation, Writing – review & editing. Vishal Win: Investigation, Writing – review & editing. Isabellah Nikitah: Investigation, Writing – review & editing. Brianna Bullock: Investigation, Writing – review & editing. Damaris Chacon: Investigation, Writing – review & editing. Lucy Sparks Mendez: Investigation, Writing – review & editing. Keita Fogus: Investigation, Writing – review & editing. Kellie Tamashiro: Resources, Funding acquisition, Writing – review & editing. Patrese A. Robinson: Supervision, Resources, Funding acquisition, Writing – review & editing. Maya Opendak: Conceptualization, Formal analysis, Supervision, Resources, Funding acquisition, Writing – original draft, Writing – review & editing.

## Data availability statement

Source data and statistics for all figures and analyses are provided in Supplementary Data 1 and will be deposited at Zenodo on acceptance.

## Declaration of generative AI use

During the preparation of this work the authors used Claude (Anthropic) to assist with statistical code, text formatting and code for figure generation. The authors reviewed and edited all content and take full responsibility for the content of the publication.

## Acknowledgements

This work was supported by the National Institutes of Health through grant R01MH133456 to MO, L40HD120184-01 to PAR, NIH DK129193 to KLT, and by core support from the Intellectual and Developmental Disabilities Research Center (IDDRC) at Kennedy Krieger Institute and Johns Hopkins University (P50HD105328; U54HD079123). Additional support was provided by the Kennedy Krieger Institute Goldstein Award, Johns Hopkins University Undergraduate Research Award (AS), the Johns Hopkins University Discovery Award (MO), the Haverford College KINSC Summer Scholars Program (BPF), and the Vivian Thomas Scholars Initiative at Johns Hopkins University (BCB). Figures were created using Python code and BioRender.

