## Supplementary Material for "Adversity severity shapes caregiving predictors and neural correlates of infant social behavior"

**Su et al.**

**Contents:**

**Supplementary Methods 1–9
Supplementary Analyses 1–8
Supplementary Figures S1–S6
Supplementary Tables S1–S2
Appendix: Maternal behavior ethogram
Supplementary Data 1: Source data and statistics workbook (separate file)**

**Supplementary Methods**

**Supplementary Methods 1. Inference with small numbers of litters**

For exact permutation tests, the reported p-value is the proportion of relabelings yielding an association at least as extreme as the observed association. Because litter means of scan proportions are multiples of 1/30 and can differ between litters only by floating-point error, litter-level values were rounded to nine decimal places before ranking so that genuine ties received tied ranks. For pup-level statistics involving a litter-level predictor, whole litters were permuted so that within-litter structure was preserved. Where clustering was accounted for by cluster-robust covariance, degrees of freedom were set to the number of litters minus one rather than the residual degrees of freedom, and estimates were confirmed by a Rademacher wild cluster bootstrap. Design effects were computed as 1 + (m − 1) × ICC to quantify the effective sample size of pup-level analyses. For the binary approach outcome, design effects were 4.08 in the full cohort (215 pups, mean 8.96 per litter, ICC = 0.387) and 1.47 in the c-Fos cohort (48 pups, two per litter, ICC = 0.471). Primary within-condition associations were additionally evaluated using exact permutation.

**Supplementary Methods 2. Maternal-behavior models**

Each of the 22 behaviors was classified as sparse if it was absent (percent duration = 0) in more than half of all litter-day recordings; by this criterion, 12 behaviors were sparse and 10 were common. Sparse behaviors were modeled as presence or absence per session using a generalized estimating equation with a binomial family, exchangeable working correlation, and litter as the clustering variable (Liang & Zeger, 1986); common behaviors were modeled using the proportion of scored bins with the same clustering structure. One model was fitted per behavior. For each behavior, the significance of rearing condition, day, and condition × day terms was assessed by Wald tests on the corresponding parameter subsets. Benjamini–Hochberg false-discovery-rate correction (Benjamini & Hochberg, 1995) was applied to the condition × day interaction across the 20 behaviors whose fits were interpretable (Table S2).

**Supplementary Methods 3. Transition entropy**

Because this analysis required sequence-resolved event records rather than the binned time-budget summaries used for the primary behavioral measures, it was conducted on the subset of sessions for which these records were available, following the information-theoretic approach used to characterize maternal-signal unpredictability in this literature (Davis et al., 2017; Vegetabile et al., 2019; Supplementary Methods). Maternal behavior onsets were ordered in time to yield a behavioral sequence, from which a first-order transition probability matrix and the empirical stationary distribution of behavioral states were estimated. The entropy rate was computed with the plug-in estimator as H = −Σᵢ π(i) Σⱼ P(i,j) log₂ P(i,j), where P(i,j) is the probability of transitioning from behavior i to behavior j and π(i) is the stationary probability of state i. The Miller–Madow bias correction was applied as a sensitivity analysis, as were restriction to maternal behaviors and use of a fixed common state space.

Entropy rate was computed per session and averaged across each litter’s available sessions (93 sessions across 21 litters, seven per rearing condition). Because sessions differed in the number of distinct behaviors recorded, and entropy computed on a session-specific state space tracks that count, this measure indexes repertoire breadth as well as sequential structure. Given the size of this subset, uneven day-level sampling, and this confound, the analysis was treated as exploratory.

**Supplementary Methods 4. Litter clustering of approach**

The proportional-hazards assumption for the condition log-rank comparisons was checked by regressing Schoenfeld residuals on transformed time and was not rejected (global *p =* 0.80); hazard ratios were treated as secondary because condition differences were concentrated in the proportion of pups approaching rather than in latency among pups that approached. To quantify the degree to which approach was structured by litter within each rearing condition, one-way random-effects intraclass correlation coefficients (ICC(1)) were computed on log10 approach latency, with nonapproaching pups entered at the 1800-s ceiling, separately by condition and for the full behavioral and neural cohorts. Ninety-five percent confidence intervals were obtained by bootstrap resampling of litters (4,000 resamples). The contrast between severe scarcity and the two milder conditions was tested by permuting condition labels across litters (5,000 permutations, one-sided). Because clustering within severe scarcity was strong in both cohorts, litter-level summaries were used for all within-condition analyses, including analyses in the neural cohort. The pup-level condition contrast in the neural cohort was retained, with within-litter dependence considered in interpreting the result.

**Supplementary Methods 5. Pup-level Cox models**

As a pup-level complement to the litter-level caregiving correlations, Cox proportional-hazards models were fitted to all 215 pups with the litter-level caregiving predictor (z-scored), rearing condition, sex, and a predictor × severe-scarcity interaction. Because pups were clustered within litters, the interaction was tested by permuting condition labels across litters (2,000 permutations), and its confidence interval was obtained by bootstrap resampling of litters (1,000 resamples).

**Supplementary Methods 6. Zone-occupancy phenotypes**

Zone-occupancy phenotypes were derived descriptively by k-means clustering of each pup’s eight-zone occupancy profile (shares of session time, square-root transformed). k = 6 was selected based on silhouette and interpretability, and each pup was labeled according to the zone in which the greatest proportion of the session was spent. The relation of each pup’s dominant zone to the first zone it entered was examined using first-entry latencies, and the share of the remaining session spent in the dominant zone after first entry was computed for each approaching pup (Supplementary Analysis 1; Fig. S6).

**Supplementary Methods 7. c-Fos quantification and pup selection**

Color thresholds in ImageJ were centered on hue 100–255 and brightness 0–140, with the lower hue and upper brightness thresholds adjusted by up to ±10 and ±15 units, respectively, when necessary to accommodate section-to-section variation in staining intensity and background; all adjustments were made without knowledge of experimental condition. The “Open” and “Watershed” functions were then applied to reduce background noise and improve particle separation before particles of 10–150 μm² were counted.

**Supplementary Methods 8. Regional c-Fos analyses**

Each *a priori* behavioral composite was computed from rank-transformed zone values across the three reachable contact zones: an approach-speed composite from the three zone latencies and a contact-time composite from the three zone occupancies. Because condition × c-Fos interaction estimates were sensitive to whether densities were entered on the raw, log, or rank scale, condition dependence was described from the within-condition estimates rather than inferred from the interaction. Pup transport and dITC c-Fos were entered together in a model of the primary outcome to assess their joint association with approach, and robustness was evaluated by leave-one-litter-out resampling. To determine whether the c-Fos association was also present within litters, the two imaged pups from each litter were contrasted (faster- minus slower-approaching sibling, log density) and tested by Wilcoxon signed-rank. This comparison removes litter-level influences shared by the sibling pair, including measured litter-level caregiving. One additional analysis was conducted as a defined multiple-comparison family. Because the LHb has been implicated in the development of caregiver-directed social behavior, the seven organized nurturing behaviors quantified here were tested against LHb c-Fos using percent duration, with Benjamini–Hochberg correction across the seven tests. A wider screen of all maternal behaviors against all four regions using both behavioral measures was exploratory and is reported in Supplementary Analysis 7.

**Supplementary Methods 9. Sex analyses**

For the transport-approach and milk ejection-LHb relationships, associations were computed separately in female and male pups; sex differences were tested using Steiger’s test for dependent correlations and a sex × predictor interaction rather than by comparing statistical significance within each sex. Because littermates are not independent, sex differences in pup-level c-Fos were tested within litters by comparing the female and male pup from the same litter using Wilcoxon signed-rank tests; unpaired comparisons across all pups were not reported.

**Supplementary Analyses**

**Supplementary Analysis 1. Behavioral phenotype: cohort, clustering, and censoring sensitivity**

In the full behavioral cohort, 78.9% of severe-scarcity pups approached the dam (56 of 71), compared with 94.4% under moderate restriction (68 of 72) and 83.3% under normative rearing (60 of 72). Because pups were clustered within litters, condition differences were evaluated using litter-aware inference. The severe-scarcity versus moderate-restriction contrast was not significant by GEE with litter clustering (*p =* 0.071) or by permutation of litter-level approach proportions (*p =* 0.24), and the full-cohort survival comparison was not significant by whole-litter permutation (*p =* 0.38). The three-condition omnibus test was also not significant (*p =* 0.071). In the balanced neural subset, in which each litter contributed two pups (design effect = 1.47), severe scarcity differed from the two milder conditions combined (log-rank χ² = 4.32, *p =* 0.038), although the three-condition omnibus test was not significant (*p =* 0.107).

The condition pattern primarily reflected the occurrence of non-approach rather than latency among pups that approached. Among approachers, latency did not differ by condition in the neural subset (*p =* 0.57) or the full cohort (*p =* 0.66). Non-approach occurred in 18.8%, 0%, and 6.2% of pups in the neural subset and 21.1%, 5.6%, and 16.7% in the full cohort for severe scarcity, moderate restriction, and normative rearing, respectively. Thus, evidence for reduced approach under severe scarcity was present in the balanced neural cohort but was weaker in the full behavioral cohort, and the condition pattern primarily reflected whether pups approached rather than latency among pups that approached.

Zone-occupancy phenotypes. Each pup’s eight-zone occupancy profile—the proportion of the 1800-s session spent in the ventrum, face, front-tail, on-back, back-head, back-tail, behind-dam, and away zones—was clustered using k-means after square-root transformation (Supplementary Methods). Six clusters were retained (silhouette = 0.52). Each cluster was dominated by a single zone, with the mean proportion of the session spent in the dominant zone ranging from 0.68 to 0.97; pups were therefore labeled according to their dominant zone (Fig. S6A).

Dominant-zone occupancy did not simply reflect the first zone encountered. Among the 184 pups that approached, only 36% ultimately became dominant in the zone they entered first, and 90 of 184 pups entered six or seven of the seven contact zones. Ventrum-dominant pups also typically entered the face or front-tail zone first. Once pups entered their eventual dominant zone, however, they spent a median of 84% of the remaining session there; 89% of approachers spent more than half of the remaining session in that zone. This proportion did not differ by rearing condition.

Dominant-zone phenotype was structured by litter (Cramér’s V for litter × phenotype = 0.54, p < 0.001 by permutation of pups across litters within condition). This litter structure was detected in each rearing condition (V = 0.56, 0.40, and 0.53; all p ≤ 0.006) and did not differ detectably between severe scarcity and the two milder conditions (ΔV = +0.05, litter-permutation *p =* 0.15). Among pups that approached, 43% of severe-scarcity pups were ventrum-dominant, compared with 28% under moderate restriction and 13% under normative rearing. No severe-scarcity pup was back-tail-dominant (0 of 71, compared with 10 of 72 under moderate restriction and 14 of 72 under normative rearing). At the litter level, the proportion of ventrum-dominant pups differed across conditions with Kruskal–Wallis *p =* 0.09, whereas the proportion dominant in any nipple-contact zone did not differ (*p =* 0.43; Fig. S6B–D).

Within severe scarcity, the proportion of ventrum-dominant approachers was not associated with typical transport (ρ = −0.33, exact *p =* 0.46) or rough transport share (ρ = +0.31, *p =* 0.50). Regional c-Fos in the neural cohort also did not differ by dominant-zone phenotype (all Kruskal–Wallis p ≥ 0.19; ventrum- versus face-dominant approachers, all p ≥ 0.20). Zone-occupancy phenotype therefore provides an additional spatial description of pup behavior during the approach test but was not detectably related to the within-scarcity caregiving or neural associations. Because ventrum occupancy could reflect nipple-directed motivation, thermotaxis toward the warmest position on the anesthetized dam, or other features of dam contact, and nipple attachment was not scored, we do not assign a specific functional interpretation to this phenotype.

**Supplementary Analysis 2. Secondary maternal-behavior comparisons**

Maternal stepping showed the clearest change associated with resource restriction. The probability that a severe-scarcity litter showed stepping increased from 0.38 to 1.00 across the exposure period, whereas it remained low under normative rearing, at 0.25 on the first day and 0.12 on the last. Stepping increased with restriction severity across all three measures (Spearman ρ = 0.60 for occurrence, 0.58 for percent duration, and 0.61 for scan proportion; all p ≤ 0.003). In contrast, nutritive nursing, arch-back nursing, side nursing, milk ejection, pup grooming, typical pup transport, total nursing duration, and nursing-posture composition did not differ across conditions (all p ≥ 0.19). Rough transport and time in the nest showed uncorrected condition differences on scan proportion, but neither survived false-discovery-rate correction across the behavior panel (smallest *q =*  0.20).

Maternal cage biting, a stress-related behavior, was recorded in 17 of 117 sessions and in 11 of 24 litters, with litter-level rates approximately twice as high under restriction as under normative rearing (0.18 under severe scarcity and 0.18 under moderate restriction versus 0.08 under normative rearing). This difference was not supported by tests treating litter as the unit of inference (Kruskal–Wallis on litter rates *p =* 0.39; whole-litter permutation *p =* 0.43; restricted conditions combined versus normative rearing *p =* 0.19). The generalized estimating equation for cage biting reported in Table S2 showed separation because no cage biting occurred under normative rearing during the first three days of the exposure window, producing coefficients corresponding to odds ratios in the thousands. These nominal p-values were therefore not interpreted, and cage biting was excluded from the false-discovery-rate correction and is reported for completeness.

**Supplementary Analysis 3. Exploratory transition entropy**

Because this analysis required sequence-resolved event records rather than the binned time-budget summaries used for the primary behavioral measures, it was conducted on the subset of sessions for which these records were available, following the information-theoretic approach used to characterize maternal-signal unpredictability (Davis et al., 2017; Vegetabile et al., 2019). Maternal behavior onsets were ordered in time to yield a behavioral sequence, from which a first-order transition matrix and empirical stationary distribution were estimated. Entropy rate was computed using the plug-in estimator. Miller–Madow correction, restriction to maternal behaviors, and use of a fixed common state space were examined as sensitivity analyses. Entropy was computed for each session and averaged across each litter’s available sessions (93 sessions across 21 litters; seven litters per condition).

Transition entropy did not differ across rearing conditions (Kruskal–Wallis *p =* 0.28; means 1.71, 1.67, and 1.54 bits) and was not associated with the percentage of pups approaching (ρ = −0.001, *p =* 0.995) or log geometric-mean approach latency (ρ = −0.16, *p =* 0.48). Sensitivity analyses were likewise null (all p ≥ 0.18). Entropy was strongly associated with the number of distinct behaviors represented in a session (ρ = +0.73), indicating that in this dataset the measure substantially indexed repertoire breadth as well as sequential structure. The absence of an association with rearing condition or social approach therefore should not be interpreted as evidence against caregiving unpredictability as a developmental mechanism.

**Supplementary Analysis 4. Robustness and boundary analyses for typical pup transport**

Amount of typical transport. Typical pup transport did not differ across rearing conditions (Kruskal–Wallis *p =* 0.67 for scan proportion; *p =* 0.99 for percent duration). Within severe scarcity, its association with approach was directionally stable in leave-one-litter-out analyses, although individual observations had substantial leverage with eight litters. Across all 24 litters, the association was weaker (ρ = −0.47, *p =* 0.020) and was substantially reduced when severe-scarcity litters were removed (ρ = −0.16, *p =* 0.54). Day-resolved features of typical transport, including linear slope across the five recording days, day-to-day standard deviation, and early- and late-window means, did not predict approach beyond the litter mean; the standard deviation and window means were themselves strongly correlated with the litter mean.

Formal transport × condition interactions were not significant for the amount of typical transport. Typical transport spanned 0–0.40 under severe scarcity but only 0–0.10 under normative rearing, limiting comparison of the association across conditions. The data therefore support a within-scarcity association for typical-transport amount but not statistical moderation by rearing condition. The association was not readily attributable to differences in outcome variance: litter-level approach outcomes showed comparable variability under normative rearing and severe scarcity (SD 450 versus 613 s; Levene *p =* 0.62), while no transport–approach association was detected under normative rearing. In addition, the exact permutation test conditions on the observed marginal distributions, preserving any compression in the observed sample under the null.

Composition of transport. Transport composition showed a different pattern. The severe-scarcity × rough-share interaction was significant on the rank scale (t = +3.00, *p =* 0.007), across every combination of raw and rank predictor scaling with raw, rank, and log outcome scaling (*p =* 0.004–0.013), and in all 23 leave-one-litter-out subsets (maximum *p =* 0.040). Within-condition correlations were ρ = +0.75 under severe scarcity, −0.04 under moderate restriction, and −0.32 under normative rearing (Fig. 3B). Unlike typical-transport amount, rough share spanned a similar and broad range across conditions (0.20–1.00, 0.25–1.00, and 0.00–1.00, respectively), permitting a more interpretable test of condition moderation. Because the interaction was estimated from eight litters per condition and was not specified in advance, it is treated as exploratory.

A pup-level Cox proportional-hazards model including all 215 pups (Supplementary Methods) provided convergent evidence. With rough share z-scored and rearing condition, sex, and a rough-share × severe-scarcity term included in the model, the interaction log-hazard ratio was −1.20 (litter-bootstrap 95% CI −2.52 to −0.21; litter-permutation *p =* 0.016). Within severe scarcity, the hazard ratio was 0.42 per SD increase in rough share (95% CI 0.14–0.81), compared with 1.22 outside severe scarcity. The corresponding model for typical-transport amount yielded a within-scarcity hazard ratio of 1.64 per SD, but the interaction was not significant (log-hazard ratio +0.40, 95% CI −0.16 to +1.48, *p =* 0.20), consistent with the litter-level analysis.

Specificity and measurement sensitivity. The within-scarcity association was specific to typical rather than rough transport when transport was considered as an amount. Rough transport did not predict approach within severe scarcity (ρ = −0.23, *p =* 0.59) and added no explanatory value when modeled with typical transport. Rough share was nearly collinear with typical transport within severe scarcity (ρ = −0.82), indicating that the two measures were complementary rather than independent descriptions of transport organization. Total transport predicted approach within severe scarcity when measured by scan proportion (ρ = −0.77, exact *p =* 0.032) or bout counts (ρ = −0.91, exact *p =* 0.005), but not percent duration (ρ = −0.33, *p =* 0.43). Total transport also showed a condition-related trend (Kruskal–Wallis *p =* 0.098), reflecting a trend in its rough component (*p =* 0.028 uncorrected, not surviving correction across the behavior panel), and therefore did not satisfy the preserved-behavior premise motivating the within-scarcity analysis. After mutual adjustment, neither typical nor total transport retained a clear independent association with approach (typical given total, partial ρ = −0.37; total given typical, partial ρ = −0.44), as expected for measures that share most of their variance. Sex-stratified estimates were similar in direction and magnitude (female ρ = −0.80; male ρ = −0.68). Because transport bouts were brief (median under 2 s), bout counts and scan proportion provided better resolution than percent duration; scan proportion was therefore retained as the primary measure and bout counts as confirmation.

**Supplementary Analysis 5. Regional specificity and condition sensitivity of the intercalated-cell association**

Within severe scarcity, dITC c-Fos showed the strongest regional association with censored approach latency (ρ = −0.88, 95% CI [−1.00, −0.39]), followed by vITC, BLA, and LHb (ρ = −0.71, −0.50, and −0.33, respectively). Convergent behavioral composites showed the same regional ordering (approach-speed composite: dITC ρ = +0.81, 95% CI [+0.23, +1.00]; contact-time composite: dITC ρ = +0.76, 95% CI [+0.06, +1.00]). Exact permutation supported the dITC association with primary approach latency (*p =* 0.007), approach speed (*p =* 0.022), and contact time (*p =* 0.037).

Evidence for condition dependence and regional specificity was weaker. Formal dITC × condition interactions were not significant on the prespecified log-density scale (primary outcome *p =* 0.22; approach-speed composite *p =* 0.065) and varied with raw, log, or rank specification. c-Fos measures were also strongly correlated across amygdala subregions (ρ = 0.68–0.80), and no individual region survived correction across all region-by-outcome tests (dITC *q =*  0.15). Within severe scarcity, the dITC association remained after adjustment for BLA activity (partial ρ = −0.84), whereas the BLA association was attenuated after adjustment for dITC (partial ρ = +0.06). These analyses identify dITC as the leading regional correlate within severe scarcity but do not establish anatomical specificity or formal moderation by rearing condition. Correlations among regional c-Fos densities did not differ detectably by rearing condition (BLA–dITC ρ = 0.69, 0.76, and 0.73 under severe scarcity, moderate restriction, and normative rearing; all severe-scarcity-versus-other-condition differences p ≥ 0.43 by litter permutation).

Within-litter contrasts likewise did not reproduce the dITC–approach association. The faster-approaching sibling showed higher dITC c-Fos in 11 of 23 pairs (median difference −0.16 log10; Wilcoxon *p =* 0.45), including 3 of 7 pairs under severe scarcity; the eighth scarcity pair was uninformative because neither imaged pup approached (both censored at 1800 s). Corresponding comparisons were also not significant for BLA (7 of 23 pairs, *p =* 0.13) or LHb (9 of 23 pairs, *p =* 0.23). Thus, the dITC–approach association was evident between litters but was not detected within littermate pairs. The within-pair comparison was informative: siblings differed in time spent on the ventrum by more than 10 percentage points of the test in 8 of 24 pairs (median absolute difference 2.9 points, interquartile range 0.1–19.2), yet the sibling with more ventrum contact had lower dITC c-Fos in only 10 of the 20 discordant pairs (sign test *p =* 1.0; Spearman ρ of paired differences = +0.12, *p =* 0.63), indicating that the litter-level dITC–approach association is not explained by the amount of maternal contact experienced during the test itself. One exploratory exception was observed for vITC: the faster-approaching sibling showed lower vITC c-Fos in 15 of 19 pairs (sign test *p =* 0.02; Wilcoxon *p =* 0.12), a pattern opposite in direction to the litter-level vITC association and not interpreted further.

Across all litters, maternal stepping was negatively associated with dITC c-Fos (ρ = −0.40, one-tailed permutation *p =* 0.027; 100,000 label permutations), and the association remained after adjustment for rearing condition (partial ρ = −0.59, one-tailed *p =* 0.002 from 20,000 within-condition permutations). Stepping itself was not associated with approach (ρ = +0.07, *p =* 0.73) and did not account for the dITC–approach relationship.

**Supplementary Analysis 6. Typical transport and dITC activity as separable correlates**

Typical transport and dITC c-Fos were not detectably associated across all 24 litters (ρ = −0.08, *p =* 0.72) or within severe scarcity (ρ = +0.61, exact *p =* 0.12; Fig. S5). Joint models were limited by the eight severe-scarcity litters and did not provide evidence that dITC recruitment accounted for the transport–approach association. Rough transport share showed a corresponding association with vITC c-Fos within severe scarcity (ρ = −0.74, exact *p =* 0.042), but this association did not survive correction across the four regions (*q =*  0.16), and neither transport measure retained an independent association after adjustment for the other. Substituting litter-level RMST across all tested pups for the imaged-pup approach summary weakened but did not eliminate the dITC–approach association (ρ = −0.76, exact *p =*  0.037). These analyses provide no evidence for a transport → dITC → approach pathway and support treating maternal transport and intercalated-cell recruitment as separable correlates of social approach.

**Supplementary Analysis 7. Robustness and sex analyses for the milk-ejection–LHb association**

Greater maternal milk ejection was associated with lower LHb c-Fos (standardized β = −0.54, Spearman ρ = −0.56, *p =* 0.005; exact permutation *p =* 0.005). Within the defined family of seven organized nurturing behaviors tested against LHb c-Fos, milk ejection was the only association surviving correction (*q =*  0.032). The remaining behaviors were nutritive nursing (ρ = +0.26, *p =* 0.22), pup grooming (ρ = −0.11, *p =* 0.61), typical pup transport (ρ = +0.08, *p =* 0.70), arch-back nursing (ρ = −0.08, *p =* 0.71), side nursing (ρ = −0.06, *p =* 0.77), and anogenital licking (ρ = −0.04, *p =* 0.85; all q ≥ 0.75). The milk-ejection association was supported by litter-resampled confidence intervals (95% CI [−0.79, −0.20]) and remained after adjustment for rearing condition (β = −0.57, *p =* 0.004). Neither milk ejection nor LHb c-Fos differed by rearing condition.

The association was sensitive to how milk ejection was quantified. Milk ejection measured as scan proportion was not associated with LHb c-Fos (ρ = −0.17, *p =* 0.43), although scan proportion and percent duration were themselves correlated (ρ = +0.61). Percent duration reflected both the number of ejections per session and their length (litter-level ρ = 0.56 and 0.61 with duration; ρ = 0.05 with each other, 22 litters with bout counts); the LHb association was carried by duration rather than count (duration adjusted for count, partial ρ = −0.46, *p =* 0.03; count adjusted for duration, ρ = +0.13, *p =* 0.56). In the wider exploratory screen of all maternal behaviors against all four regions using both behavioral measures (128 tests), milk ejection–LHb was the strongest association, but no individual test survived correction across this broader family (*q =*  0.55). The milk-ejection–LHb relationship is therefore supported within the defined nurturing-behavior family but requires independent confirmation.

LHb c-Fos was not associated with pup social approach (geometric-mean latency of the imaged pups ρ = −0.18, *p =* 0.40; litter RMST across all tested pups ρ = −0.35, *p =* 0.09; both across 24 litters). Female pups showed higher LHb c-Fos than their male littermates in 14 of the 20 litters contributing a female–male pair (geometric means 117 versus 64 cells/mm²), but the paired difference was not significant (Wilcoxon signed-rank *p =* 0.070, *q =*  0.25; sign-flip permutation *p =* 0.091), and no region differed by sex in paired analyses (all q ≥ 0.25). The milk-ejection association was not detectably modified by sex: milk ejection remained significant in models accounting for sex, and neither the sex × predictor interaction nor Steiger’s test supported a sex difference in the association (*p =* 0.17 and *p =* 0.11, respectively).

**Supplementary Analysis 8. Corticosterone sensitivity analyses**

Plasma corticosterone was measured by ELISA in P12 littermates from 12 litters (four per condition) and analyzed at the litter level. Concentrations followed a non-monotonic condition pattern (Kruskal–Wallis *p =* 0.044; means 4.26, 7.06, and 3.34 ng/mL for severe scarcity, moderate restriction, and normative rearing). The omnibus effect was carried by moderate restriction versus normative rearing (*p =* 0.029); severe scarcity was directionally elevated relative to normative rearing but did not differ significantly (*p =* 0.49).

Corticosterone was not significantly associated with typical pup transport (ρ = +0.41, *p =* 0.19), and leave-one-litter-out estimates remained non-significant. No regional c-Fos measure was significantly associated with corticosterone, and all litter-resampled confidence intervals included zero (Fig. S3). Corticosterone also did not differ by pup sex (Wilcoxon signed-rank *p =* 0.15). These analyses did not detect covariance between P12 circulating corticosterone and the principal caregiving, neural, or behavioral relationships. Given the small sample and measurement of corticosterone in separate littermates, they do not establish mechanistic independence from stress-hormone signaling.

**Supplementary Figures**

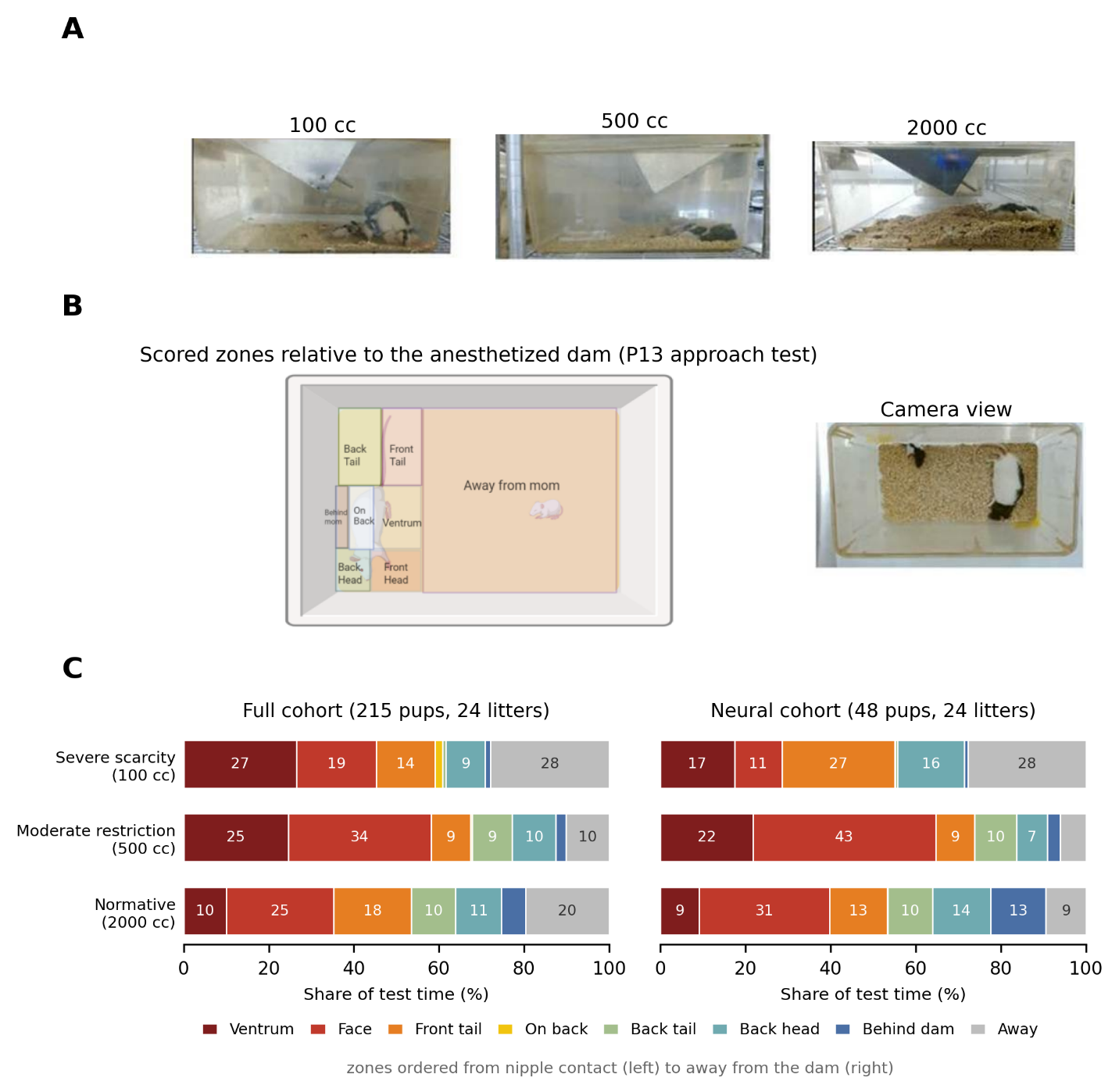

**Figure S1. Experimental apparatus and zone occupancy.** (A) Representative photographs of the bedding manipulation conditions illustrating relative bedding volume across severe scarcity (100 cc), moderate restriction (500 cc), and normative rearing (2000 cc). (B) The P13 pup–dam approach apparatus, showing the spatial zones scored relative to the anesthetized dam, from the direct nipple-contact zones (ventrum, face, front tail) to the zone away from the dam, and a representative image of the standardized camera view used for behavioral quantification. (C) Share of test time spent in each zone by rearing condition (condition mean of pup-level occupancy, rescaled to sum to 100%; no individual pup divides its time this way, because each session is dominated by a single zone, see Fig. S6), for the full behavioral cohort (215 pups, 24 litters) and the neural cohort (48 pups, two per litter); zones are ordered from nipple contact to away from the dam. Severe-scarcity pups spent the largest share of the test away from the dam in both cohorts, consistent with their reduced approach (Fig. 1B), although zone shares did not differ between conditions at the litter level (litter-mean time away, Kruskal–Wallis *p =* 0.90; contact-zone composite *p =* 0.77). The condition-mean profile is dominated by the same zones that define the individual-pup phenotypes in Fig. S6, and the away share in each condition largely reflects its proportion of non-approaching pups.

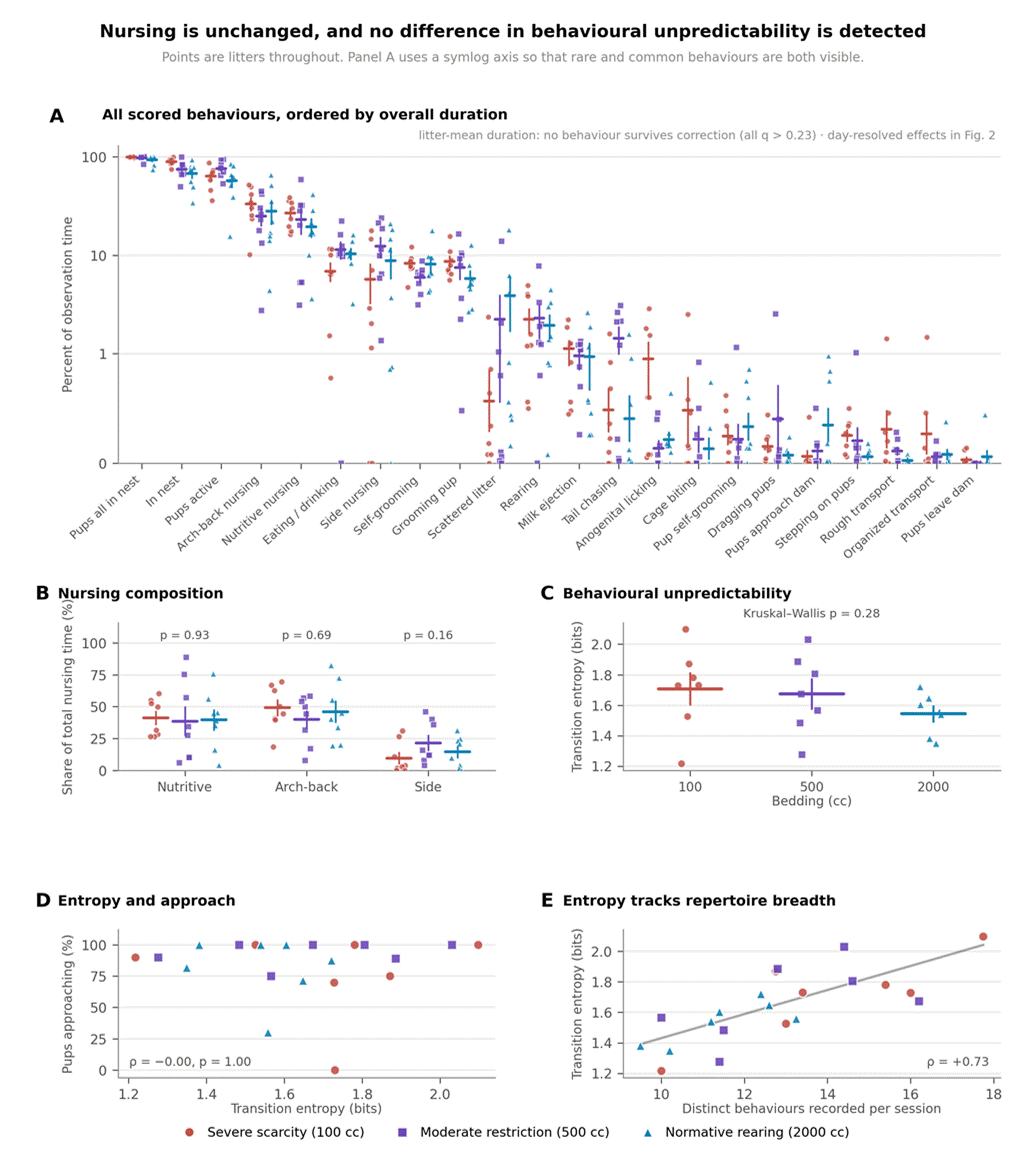

**Figure S2. Nursing quantity and composition are unchanged by rearing condition, and no difference in behavioral unpredictability is detected. (**A) Percent of observation time for all 22 scored behaviors, ordered by overall mean duration. Points are litter means (24 litters, 117 sessions); horizontal bars are condition means ± SEM. The axis is symlog (linear below 1%) so that rare and common behaviors are both visible. Kruskal-Wallis tests on litter means with Benjamini–Hochberg correction across the 22 behaviors: no behavior survives correction (smallest *q =*  0.23; largest uncorrected effects, dam in nest *p =* 0.012 and stepping on pups *p =* 0.021). Day-resolved models are in Figure 2 and Table S2. (B) Each nursing posture as a share of total nursing time. Kruskal-Wallis *p =* 0.93 (nutritive, 41/39/40%), 0.69 (arch-back, 49/40/46%), and 0.16 (side, 10/22/14%). Total nursing duration also did not differ (*p =* 0.37; 66/60/56% of observation time). (C) Transition entropy of the maternal behavior sequence, one value per litter, computed over the 93 sessions from the 21 litters with surviving event-level records (seven per condition). Kruskal-Wallis *p =* 0.28; means 1.71, 1.67, and 1.54 bits. (D) Entropy against the percentage of pups approaching (ρ = −0.001, *p =* 0.995; against log geometric-mean latency, ρ = −0.16, *p =* 0.48). (E) Entropy against the number of distinct behaviors recorded per session (ρ = +0.73; Theil–Sen fit). With 22 behaviors and sessions of this length the entropy measure largely indexes repertoire breadth, so it cannot cleanly separate sequential unpredictability from how many behaviors were logged; the null in (C) should be read accordingly.

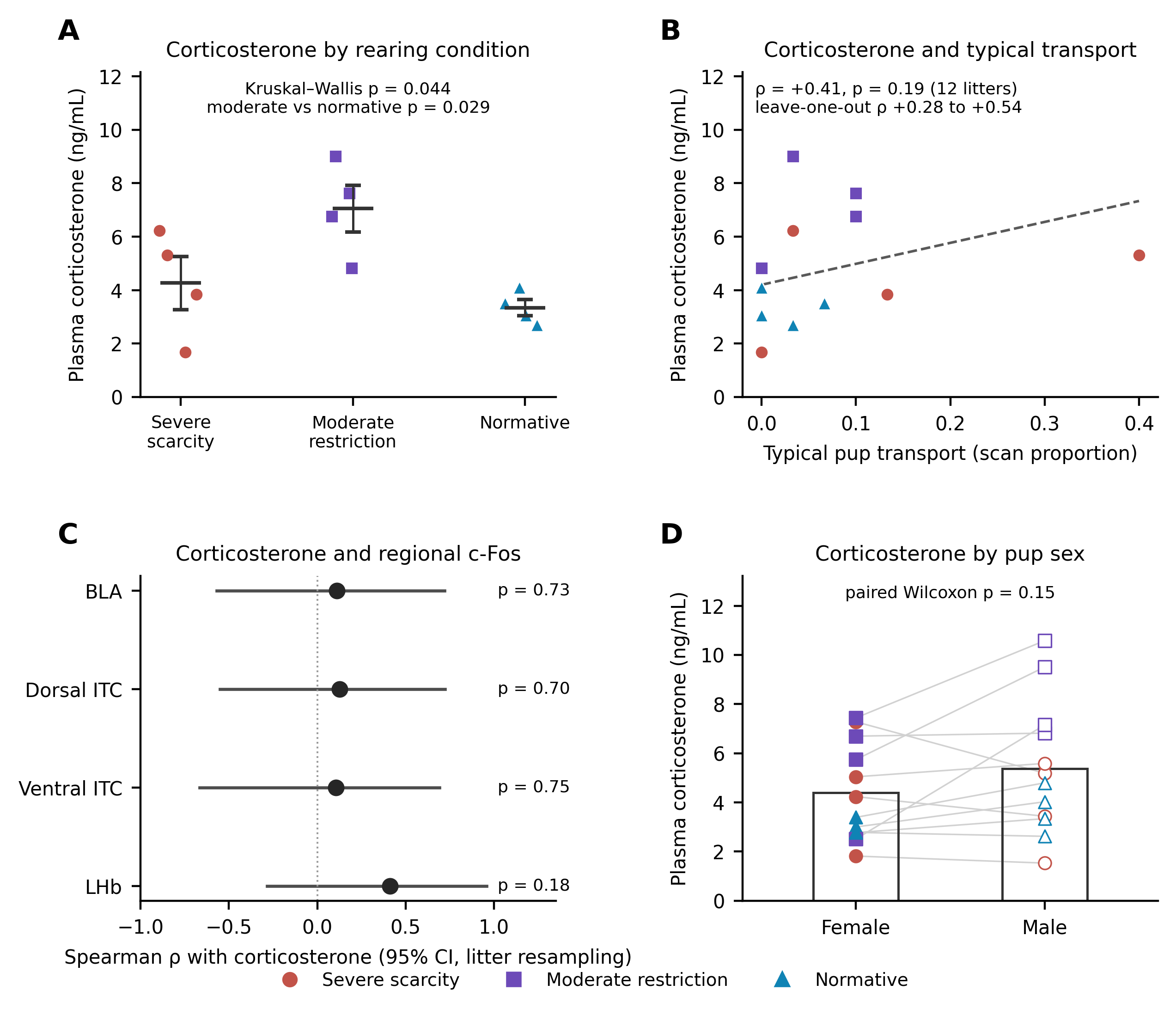

**Figure S3. Corticosterone is dissociated from caregiving, neural, and behavioral measures.** Plasma corticosterone was measured by ELISA in P12 littermates, one female and one male from each of 12 litters (four per condition), and analyzed at the litter level. (A) Corticosterone by rearing condition (points, litters; bars, condition mean ± SEM). Concentrations followed a non-monotonic pattern, highest under moderate restriction (Kruskal-Wallis *p =* 0.044; means 4.26, 7.06, and 3.34 ng/mL). The omnibus effect is carried by moderate restriction against normative rearing (*p =* 0.029); severe scarcity was directionally elevated relative to normative rearing but not significantly so (*p =* 0.49). (B) Corticosterone against typical pup transport, the litter-mean scan proportion used throughout the main text. The two were not significantly related (Spearman ρ = +0.41, *p =* 0.19). Four of the twelve litters recorded no transport and litter 26-42 spans most of the observed range; leave-one-litter-out ρ runs from +0.28 to +0.54 and reaches significance in none of the twelve fits. (C) Correlations between litter corticosterone and regional c-Fos density (points, ρ; bars, 95% CI from litter resampling). No region was significantly associated with corticosterone and every confidence interval included zero. (D) Corticosterone by pup sex (points, individual pups; lines connect within-litter female and male values; bars, sex means). Corticosterone did not differ by sex (Wilcoxon signed-rank *p =* 0.15).

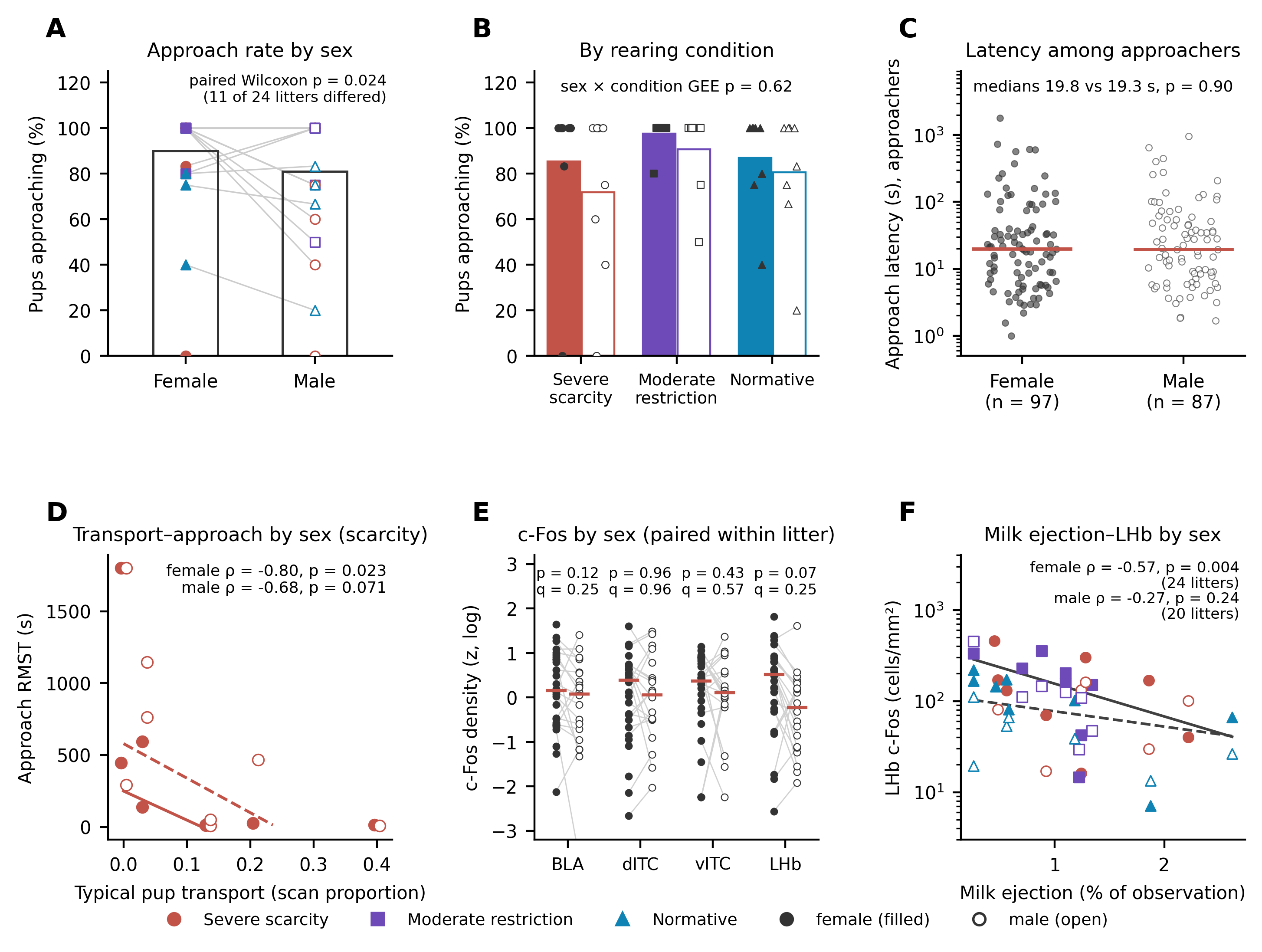

**Figure S4. Pup sex shifts approach rate, but not regional c-Fos or the caregiving relationships.** Female, filled markers; male, open markers. In (D) and (F) the two sexes share the litter's x value, because typical transport and milk ejection are litter-level measures, and are displaced horizontally by a small fixed amount so that neither point obscures the other. (A) Proportion of pups approaching the dam per litter, by pup sex (points, litters; lines connect within-litter female and male values; bars, sex means; point color denotes rearing condition). Females approached somewhat more often than males (89.9% versus 81.0%; paired Wilcoxon *p =* 0.024 across the 11 litters in which the sexes differed; in the remaining 13 litters every pup of both sexes approached). (B) The same comparison within each rearing condition. The female advantage was consistent in direction across all three conditions (85.4 versus 71.9% under severe scarcity, 97.5 versus 90.6% under moderate restriction, 86.9 versus 80.6% under normative rearing), with no sex × condition interaction (GEE *p =* 0.62). (C) Approach latency among pups that approached (log scale; bars, medians). Latency did not differ by sex (female median 19.8 s, n = 97; male 19.3 s, n = 87; *p =* 0.90). (D) Typical pup transport against the litter-level approach outcome (restricted mean survival time) within severe scarcity, computed separately for female and male pups. The relationship was present and of similar magnitude in both sexes (female Spearman ρ = −0.80, exact permutation *p =* 0.023; male ρ = −0.68, *p =* 0.071; combined ρ = −0.75, *p =* 0.040). Lines are Theil–Sen fits, truncated where they would predict a negative RMST. (E) Regional c-Fos density (z-scored on log density; bars, medians; points, individual pups; thin lines connect the female and male pup of the same litter) by sex, tested by Wilcoxon signed-rank on within-litter pairs. Twenty of the 24 litters contributed a female-male pair; the remaining four contributed two females and appear as unconnected points. LHb c-Fos was higher in the female of the pair in 14 of 20 litters, but no region differed by sex after Benjamini–Hochberg correction across the four regions (LHb *p =* 0.070, *q =*  0.25; BLA *p =* 0.12, *q =*  0.25; ventral ITC *p =* 0.43, *q =*  0.57; dorsal ITC *p =* 0.96, *q =*  0.96). One pup with zero counts in both dorsal and ventral ITC is displayed using a half-minimum offset and is retained as a true zero in the rank tests. (F) Milk ejection against LHb c-Fos separately by sex (female ρ = −0.57, *p =* 0.004 across 24 litters; male ρ = −0.27, *p =* 0.24 across 20). The association runs in the same direction in both sexes.

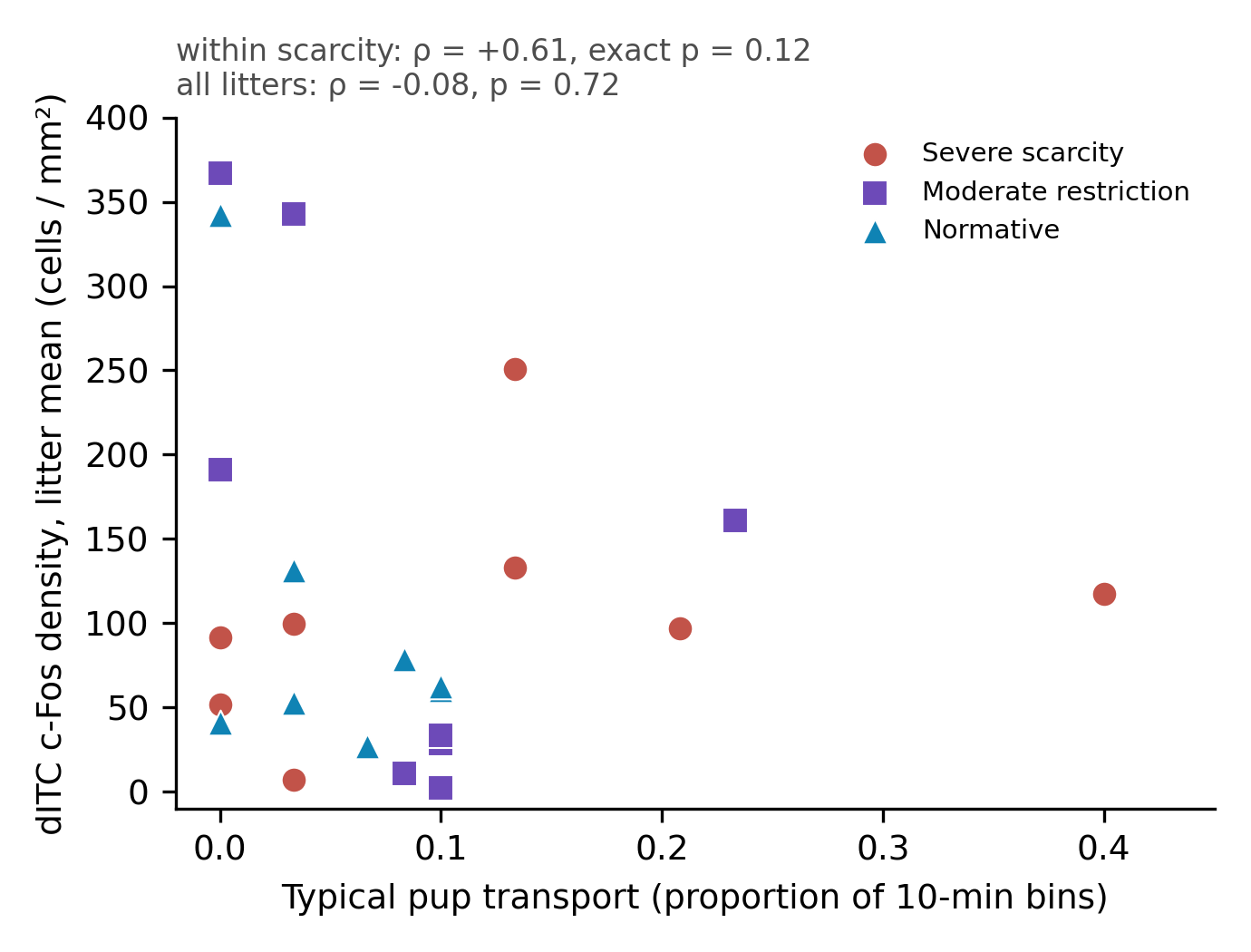

**Figure S5. Typical maternal transport and dorsal intercalated-cell c-Fos are not robustly associated.** Litter-mean dITC c-Fos density (two pups per litter) against typical pup transport (scan proportion, litter mean over scored sessions), for all 24 litters, symbols by rearing condition. Within severe scarcity the association was positive but not significant (Spearman ρ = +0.61, exact *p =* 0.12, n = 8); across all litters ρ = −0.08, *p =* 0.72. Partial correlations within severe scarcity (typical transport with approach given dITC, and dITC with approach given typical transport) are reported in Supplementary Analysis 6; neither predictor was eliminated by adjustment for the other.

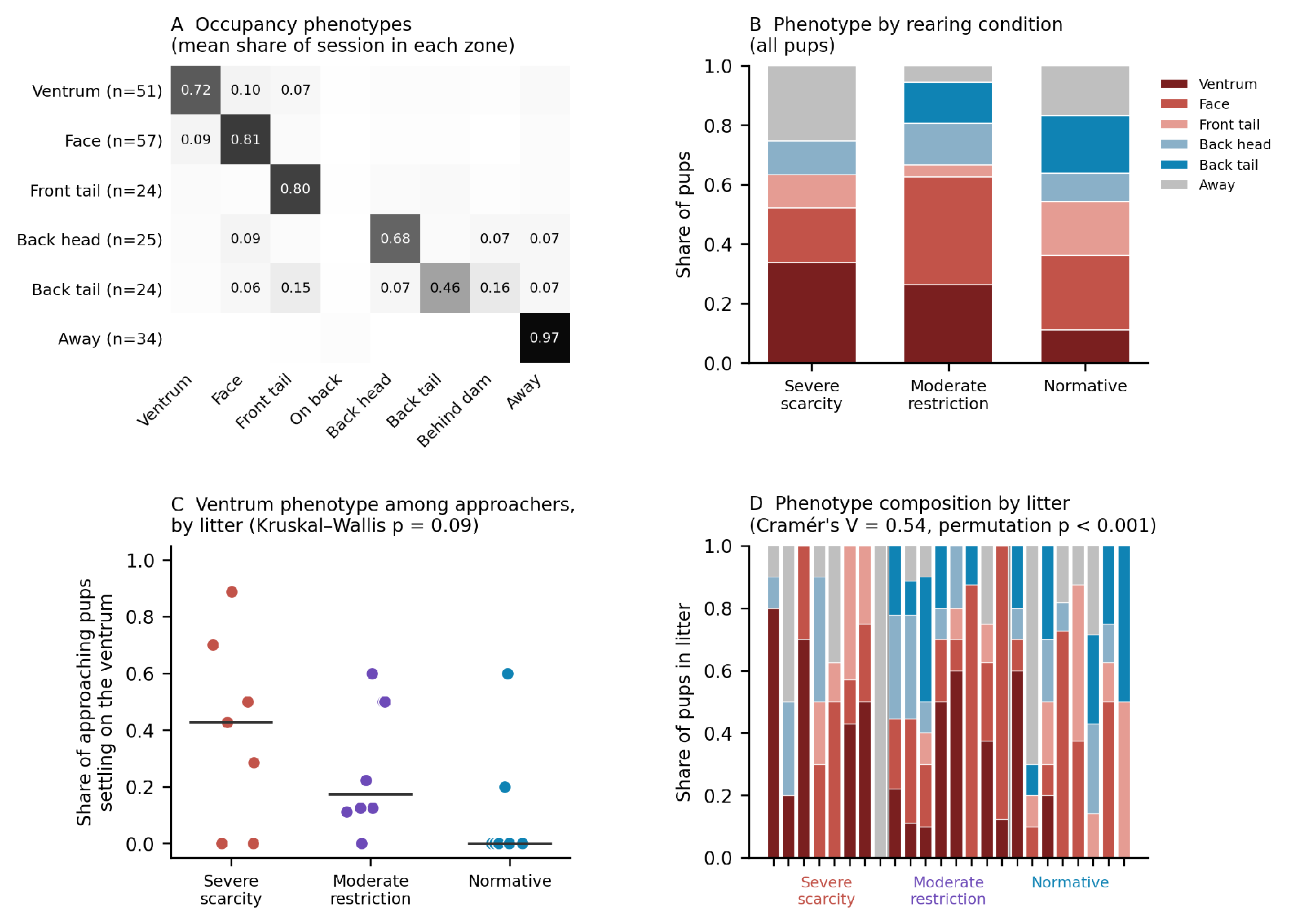

**Figure S6. Zone-occupancy phenotypes. (**A) Six phenotypes from k-means clustering of the eight-zone occupancy profile (215 pups; square-root-transformed shares of session time). Cells give the mean share of the session spent in each zone; each phenotype is dominated by one zone, and pups are labeled by that zone. (B) Phenotype composition by rearing condition, all pups. No severe-scarcity pup was back-tail-dominant (0 of 71, versus 10 of 72 under moderate restriction and 14 of 72 under normative rearing); the ventrum was the most common dominant zone under severe scarcity (24 of 71). (C) Among pups that approached, the share whose session was dominated by the ventrum, one point per litter with condition medians (0.43, 0.17, 0.00; Kruskal–Wallis *p =* 0.09). (D) Phenotype composition of each litter, grouped by condition; littermates tend to share a phenotype in every condition (Cramér's V = 0.54; p < 0.001 by permuting pups across litters within condition). Dominant zone was not associated with regional c-Fos in the neural cohort or with maternal transport within severe scarcity (Supplementary Analysis 9).

| Characteristic | Severe scarcity  (100 cc) | Moderate restriction  (500 cc) | Normative rearing  (2000 cc) | p (KW) |
| --- | --- | --- | --- | --- |
| Average pup weight at perfusion (g) | 22.9 ± 3.8 | 24.1 ± 1.8 | 26.0 ± 4.8 | 0.30 |
| Litter size | 10.1 ± 0.4 | 10.4 ± 0.7 | 10.5 ± 0.9 | 0.71 |
| Dam age at P13 (days) | 232 ± 110 | 275 ± 120 | 184 ± 67 | 0.21 |
| Dam parity (previous litters) | 1.0 ± 1.2 | 1.0 ± 1.7 | 0.4 ± 0.5 | 0.56 |

**Table S1. Litter and dam characteristics did not differ across rearing conditions.** Values are mean ± SD across litters (n = 8 per condition); pup body weight was measured at perfusion (~P13). p-values are from Kruskal–Wallis tests across the three rearing conditions. No characteristic differed significantly by condition, indicating that the behavioral, endocrine, and neural differences reported here are not attributable to differences in pup growth, litter size, or dam demographics.

| Behavior | Category | Model | Condition p | Day p | Condition × Day p | Condition × Day q (FDR) |
| --- | --- | --- | --- | --- | --- | --- |
| Step On Pups | Adverse handling | GEE (presence) | 0.174 | 0.743 | <0.001 | 0.007 * |
| Scattered Litter | Adverse handling | GEE (presence) | 0.160 | 0.151 | 0.155 | 0.515 |
| Rough Pup Transport | Adverse handling | GEE (presence) | 0.078 | 0.063 | 0.482 | 0.755 |
| Drag Pups | Adverse handling | GEE (presence) | 0.911 | 0.201 | 0.938 | 0.938 |
| Chase Tail | Maternal stress behaviors | GEE (presence) | 0.544 | 0.363 | 0.761 | 0.895 |
| Bite Cage ‡ | Maternal stress behaviors | GEE (presence) | 0.002 | 0.773 | <0.001 | — |
| Nursing On Pups | Nursing & feeding | GEE (proportion) | 0.305 | 0.592 | 0.027 | 0.270 |
| Milk Ejection | Nursing & feeding | GEE (proportion) | 0.374 | 0.229 | 0.147 | 0.515 |
| Nursing on Side | Nursing & feeding | GEE (presence) | 0.611 | 0.247 | 0.650 | 0.821 |
| Nursing Arch Back | Nursing & feeding | GEE (proportion) | 0.765 | 0.612 | 0.917 | 0.938 |
| Anogenital Licking | Nurturing contact | GEE (presence) | 0.009 | 0.948 | 0.107 | 0.515 |
| Grooming Pup | Nurturing contact | GEE (proportion) | 0.363 | 0.053 | 0.218 | 0.623 |
| In Nest | Transport & nest | GEE (proportion) | 0.584 | 0.408 | 0.049 | 0.326 |
| Pups All In Nest | Transport & nest | GEE (proportion) | 0.307 | 0.861 | 0.313 | 0.665 |
| Normal Pup Transport | Transport & nest | GEE (presence) | 0.483 | 0.476 | 0.657 | 0.821 |
| Rear | Maternal self-maintenance | GEE (proportion) | 0.557 | 0.156 | 0.324 | 0.665 |
| Self Groom | Maternal self-maintenance | GEE (proportion) | 0.278 | 0.303 | 0.333 | 0.665 |
| Drink/Eat | Maternal self-maintenance | GEE (proportion) | 0.516 | 0.241 | 0.489 | 0.755 |
| Pup Self Groom | Pup behaviors | GEE (presence) | 0.333 | 0.062 | 0.491 | 0.755 |
| Pup Approach Mom | Pup behaviors | GEE (presence) | 0.884 | 0.433 | 0.629 | 0.821 |
| Pup Active | Pup behaviors | GEE (proportion) | 0.959 | 0.301 | 0.936 | 0.938 |
| Pup Leave Mom ‡ | Pup behaviors | GEE (presence) | 0.006 | 1.000 | 0.155 | — |

**Table S2. Statistics for the 22 analyzed maternal and pup behaviors.** Each behavior was modeled across the manipulation window (postnatal day 8–12) with a generalized estimating equation using an exchangeable working correlation and litter as the clustering unit. Common behaviors (absent in at most 27% of recordings) were modeled as the proportion of scored bins in which the behavior occurred; sparse behaviors (absent in more than half of recordings, assessed on percent duration) were modeled as presence or absence with a binomial family, a session counting as containing the behavior if it was registered on either measure. Twelve behaviors were sparse and ten common; the classification is insensitive to the threshold, as any cutoff between 27% and 51% yields the same split. Behaviors are grouped by category and ordered within each category by the condition × day q-value, so that each block can be read on its own. P-values are Wald tests on the condition, day, and condition × day parameter subsets. Benjamini–Hochberg false-discovery-rate q-values are reported for the condition × day interaction across the 20 behaviors whose fits are interpretable. The two fits flagged for separation (‡) are excluded from the correction, because their nominal p-values are artifacts of the fit rather than estimates of an effect, and no q-value is reported for them. Only maternal stepping on pups survived correction (highlighted; *q =*  0.007). Maternal walking, the twenty-third behavior in the ethogram, was recorded in a single session and could not be modeled; it is omitted from the table. ‡ Fits flagged for separation: these behaviors occur in fewer than 15% of sessions, one arm of the design contains no occurrences over part of the window, and the resulting coefficients correspond to implausibly large odds ratios. Their nominal p-values are not interpreted and they are excluded from the false-discovery-rate correction; maternal cage biting is described in Supplementary Analysis 2. A separate false-discovery-rate screen relating maternal behavior to pup approach within severe scarcity is reported in the main text.

**Appendix - Maternal Behavior Ethogram**

**Introduction**

Maternal and pup behaviors were scored using a predefined ethogram adapted from established rodent maternal care paradigms. Behaviors were defined *a priori* to allow objective, reproducible scoring across observers and experimental conditions. Observations were conducted using instantaneous sampling; at each scoring timepoint, the single predominant behavior was recorded. Behaviors were mutually exclusive within a given timepoint. Definitions emphasize observable actions rather than inferred intent. Maternal behaviors were grouped into caregiving-related behaviors, nest organization, and general maternal activity, while pup behaviors were scored separately to capture pup-initiated interactions and activity.

**Maternal Care Behaviors**

**Anogenital licking**
The mother licks the pup’s anogenital region, typically while the pup is positioned ventral side up. This behavior is a core component of normative maternal care and facilitates pup elimination.

**Groom pup**
The mother grooms pups by licking body regions such as the head, torso, or limbs. This behavior reflects tactile stimulation and maternal engagement with the litter.

**Normal transporting pup**
The mother transports a pup by gently grasping the nape of the neck or upper back. This behavior reflects organized and appropriate pup handling.

**Rough transporting pup**
The mother transports a pup using an inappropriate body part (e.g., legs, abdomen, or flank). This behavior reflects atypical or disorganized maternal handling.

**Drag pup**
The mother pulls or drags one or more pups away from the nest, often while pups remain attached to the nipple. This behavior commonly occurs when the mother exits the nest following nursing or during nest disturbance.

**Mom steps on pup**
The mother steps on one or more pups, often eliciting pup vocalizations.

**Arch-back nursing**
The mother nurses while maintaining an arched posture over the pups and her front legs are rigid. This posture often coincides with milk ejection, during which pups extend their bodies simultaneously.

**Side nursing**
The mother lies on her side while pups are attached to the nipples and actively nursing.

**Nursing on pups**
The mother nurses while positioned directly on top of the pups in a blanket posture, with pups attached beneath her body.

**Milk ejection**
The mother exhibits milk ejection, identified by a pup stretch reflex characterized by pup body elongation followed by increased activity. The duration of milk ejection is recorded until pups settle.

**General Maternal Activity**

**Mom walking around**
The mother moves outside the nest area, including walking on cage or hutch structures. This category includes general locomotion and exploratory sniffing.

**Mom rearing**
The mother stands upright on her hind legs. Rearing associated with eating or drinking is not included in this category.

**Mom self-groom**
The mother grooms herself through behaviors such as licking, scratching, face wiping, or paw-assisted grooming. Body rotation during grooming may occur.

**Mom eat/drink**
The mother consumes food or water. This behavior is scored independently of locomotion or rearing.

**Bite cage**
The mother bites or gnaws on the metal cage top or walls. While brief gnawing may occur during exploration, repeated or sustained cage biting is scored as this behavior and may reflect stress-related activity.

**Chase tail**
The mother runs in tight circles while chasing and occasionally biting her tail. This behavior is considered stress-related.

**Nest Organization**

**Mom in nest**
The mother is physically located within the nest area with her pups, indicating proximity and availability for caregiving. The nest area is defined as the bedding-constructed nest formed by the dam.

**Scattered litter**
One or more pups are located outside the nest while the remaining pups are within the nest area.

**Pups all in nest**
All pups are grouped together within the nest area. This behavior is not scored if any pup is located outside the nest.

**Pup Behaviors**

**Pup approach mom**
One or more pups move toward the mother when she enters the nest or when she is positioned outside the nest area. This behavior reflects pup-initiated contact seeking.

**Pup leave mom**
One or more pups move away from the mother or disperse when she is nearby. Depending on developmental stage, this behavior may reflect avoidance or exploratory activity.

**Pup self-groom**
Pups engage in grooming behaviors such as face wiping or paw-mediated grooming. This behavior is scored only for pups younger than postnatal day 14.

**Pup(s) active/moving**
At least one pup is actively moving at the time of observation. If only a single pup is active, this is explicitly noted.
